# Fallen Pillars: The Past, Present, and Future Population Dynamics of a Rare, Specialist Coral-Algal Symbiosis

**DOI:** 10.1101/365650

**Authors:** Andrea N. Chan, Cynthia L. Lewis, Karen L. Neely, Iliana B. Baums

## Abstract

Rare and ecologically specialized species are at greater risk of extinction. Rarity in terms of low genotypic diversity may be obscured in sessile foundation species that can reproduce asexually. Asexual propagules are often only distinguishable from sexual recruits through molecular tools. Thus, molecular markers are necessary to assess genotypic variation and population structure in clonal organisms such as corals. The global decline of corals has necessitated marker development for improved conservation of rare coral species. We infer past demographic changes, describe modern population structure, and quantify asexual reproduction of the uncommon Caribbean pillar coral, *Dendrogyra cylindrus* and its endosymbiotic dinoflagellate, *Symbiodinium dendrogyrum* using *de novo* microsatellite markers. Results show that *D. cylindrus* comprises three distinct populations in the Caribbean whereas the symbiont was differentiated into four populations. Thus, barriers to gene flow differ between host and symbiont. In Florida, host and symbiont reproduced mainly asexually, yielding lower genotypic diversity than predicted from census size. Models of past demographic events revealed no evidence of historical changes in population size, consistent with the geological record of *D. cylindrus* indicating it has been rare for hundreds of thousands of years. The most recent global thermal stress event triggered a severe disease outbreak among *D. cylindrus* in Florida, resulting in a precipitous population decline. Projections indicate a high likelihood that this species may become locally extinct within the coming decades. The ecosystem consequences of losing rare coral species and their symbionts with increasingly frequent extreme warming events are not known but require urgent study.

## Introduction

Ecologically rare species are predicted to be more vulnerable to environmental change and are at greater risks of extinction with shifts in climate (Caughley, 1994; Davies et al., 2004; McKinney, 1997). Species are rare because they may inhabit narrow geographic ranges, occupy few specific habitats, and/or exhibit low abundance in nature (Rabinowitz, 1981). It is sometimes presumed that rare species tend to be competitively inferior, although work on sparse and common grasses does not support this conclusion (Rabinowitz et al., 1984). Regardless, there are obvious consequences to having low population densities, including difficulties finding a mate (Stephens & Sutherland, 1999) and vulnerability to genetic drift (Ellstrand & Elam, 1993). However, there are many rare species that are adapted to low population densities (de Lange & Norton, 2004; Flather & Sieg, 2007), and these species persist for long periods of evolutionary time.

The persistence of rare species is further challenged when they are obligate partners of specific symbionts. The endangered terrestrial orchid *Caladenia huegelii* associates with a specific mycorrhizal fungus throughout its range (Swarts et al., 2010). This specificity between partners has caused *C. huegelii* to be rare due to the limited suitable environmental conditions of the mycorrhiza. Thus, the strict niche characteristics of one partner in a symbiosis may drive the scarcity of the other. Further, intra-specific diversity in both partners and fidelity of genotype-genotype associations can play a role in how the symbiosis responds to changing conditions (Baums et al., 2014b; Parkinson et al., 2015; Parkinson & Baums, 2014).

Because sessile organisms often reproduce by asexual fragmentation, the relative rarity of these species may be obscured by an apparently large census size. Thus, molecular markers are necessary to distinguish individuals and provide a way of measuring the genotypic diversity of populations. There are clear consequences to being a clonal, rare species that is self-incompatible. Clonal populations of the rare dwarf shrub *Linnaea borealis* had low reproductive success due to a lack of nearby conspecifics, leading to increased geitonogamy (Scobie & Wilcock, 2009). Likewise, the aquatic plant *Decodon verticillatus* had reduced sexual reproduction in clonal populations at the northern range limit (Dorken & Eckert, 2001). However, extensive clonal reproduction can also be beneficial, especially for rare species, by maintaining population sizes despite a lack of sexual recruitment. Clonal reproduction was the driver of stable population size in the threatened, endemic columnar cactus *Stenocereus eruca* (Clark-Tapia et al., 2005). No sexual seedlings were observed during the study. Thus, clonal propagation can provide a “storage effect” by increasing the persistence of individual genotypes when the influx of sexual recruits is low or absent due to unfavorable environmental conditions (Boulay et al., 2014; Warner & Chesson, 1985). This storage effect is most pronounced when adults are long-lived, reproductive output is high, and population densities are low – such as in some mass-spawning corals – and thus may play an important role in maintaining intraspecies diversity (Baums et al., 2014a; Baums et al., 2006; Boulay et al., 2014).

While many marine invertebrates produce planktonic larvae and hence have great potential to disperse over large areas, species differ significantly in their abundance. Populations of rare marine species may become fragmented more easily, reducing connectivity and increasing extinction risk. Indeed populations of rare marine sponges (*Scopalina lophyropoda*) are highly structured, likely due to limited larval dispersal (Blanquer & Uriz, 2010). Despite high larval retention, populations of *S. lophyropoda* are not inbred, indicating that this species maintains genetic diversity despite low abundance. In recent coral surveys throughout the Caribbean, some species were consistently more abundant (e.g. *Orbicella faveolata*), while others were rare, such as the pillar coral *Dendrogyra cylindrus* (Edmunds et al., 1990; S. Miller et al., 2013; Steiner & Kerr, 2008; Ward et al., 2006). The fossil record supports the relative rarity of *D. cylindrus* through time, although there is localized evidence that pillar corals were more prevalent on Pleistocene reefs (Hunter & Jones, 1996). Corals in the genus *Dendrogyra* only appear 29 times in the Paleobiology Database, whereas there are 278 *Acropora palmata* fossils, 111 *Orbicella faveolata* fossils, and 160 *Meandrina* (a sister genus to *Dendrogyra*) fossils (paleobiodb.org, accessed 18 February 2018).

*D. cylindrus* forms an obligate symbiosis with another rare species, *Symbiodinium dendrogyrum* (A. Lewis et al. in review). In recent surveys of Caribbean corals, *S. dendrogyrum* (formerly ITS2 type B1k) was only found associated with *D. cylindrus*, indicating that it has a narrow habitat range (Finney et al., 2010). Increasing rarity of one species will likely lead to subsequent rarity in the other mutualistic partner. While it is unclear which partner is driving the rarity of the *D. cylindrus*-*S. dendrogyrum* mutualism, assessing levels of within-species diversity is imperative to understanding the functioning of and threats to this association. If co-dispersal between partners is occurring, then the population structure of both symbiotic species will be congruent (Werth & Scheidegger, 2012) and facilitate local adaptation (Baums et al., 2014b).

*Dendrogyra cylindrus* (Ehrenburg, 1834) is a conspicuous if uncommon Caribbean coral in the family Meandrinidae (Figure 1). It is the only species in the genus, and it occurs between 1 and 25 m depths (Goreau & Wells, 1967). This species is a broadcast spawner (Szmant, 1986) and columns of the coral colonies sometimes break apart indicating the potential for asexual reproduction via fragmentation. We used *de novo* and existing microsatellite markers for host and symbiont to assess the null hypotheses of no population structure and no asexual reproduction in either partner. We further hypothesized that coral colonies harbor *S*. *dendrogyrum* throughout the host’s range. We then used a maximum-likelihood demographic model to assess the null hypothesis of no past changes in population size in *D. cylindrus*.

**Figure 1.**
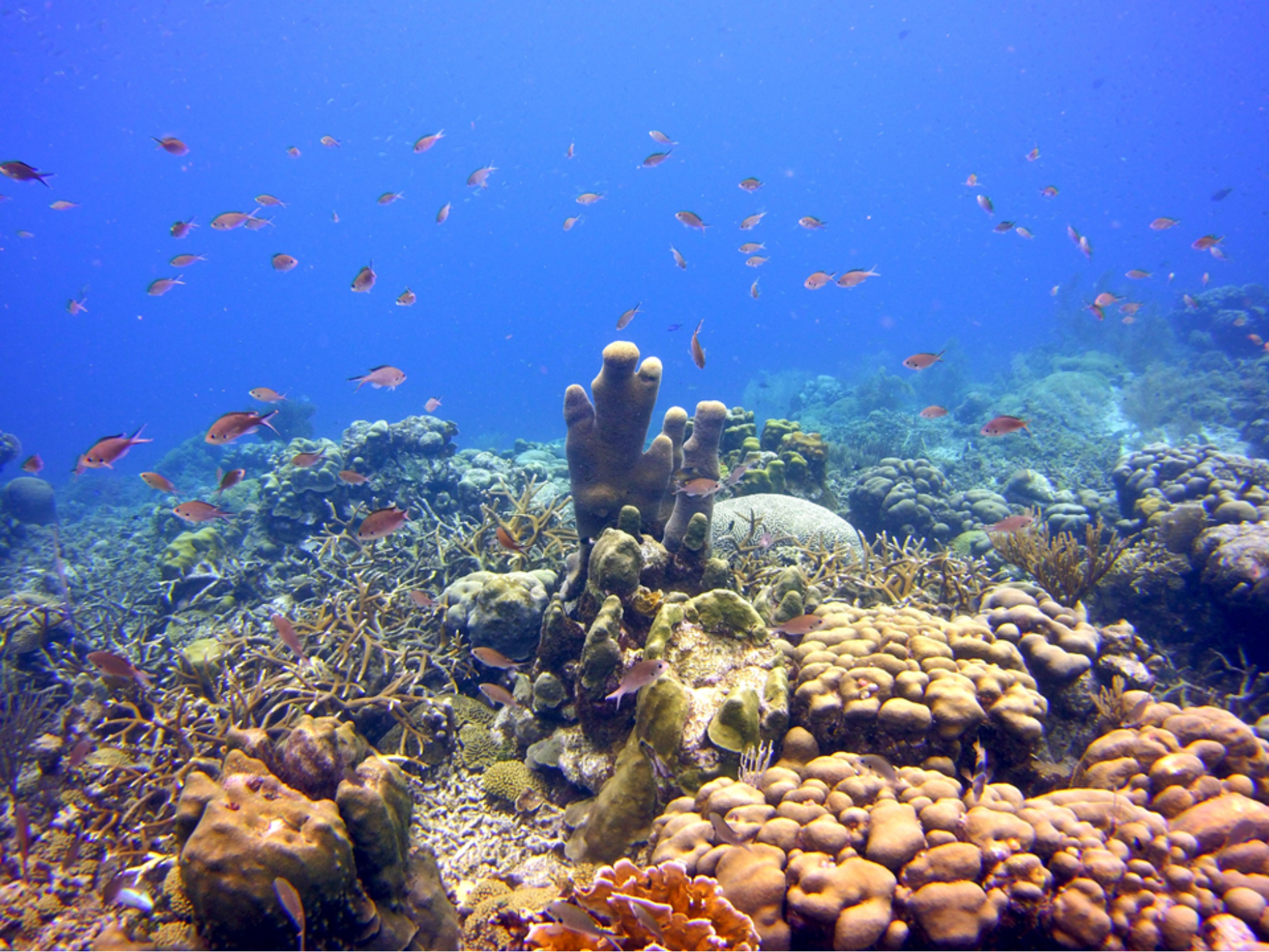
A lone colony of the pillar coral, *Dendrogyra cylindrus*, in Curaçao (center of image). Photo credit: Iliana Baums.

## Materials and Methods

### Sample Collection

Samples of *D. cylindrus* were collected from 51 sites along the Florida Reef Tract, 3 sites in Curaçao, 6 sites in the US Virgin Islands, 3 sites in Belize, and 4 sites in the Turks and Caicos Islands (Figure 2). Because of the rarity of *D. cylindrus*, we targeted sites with known occurrences of the coral instead of attempting a random sampling scheme. Sampling intensity was higher in Florida, where multiple colonies were sampled from single sites. Also, within-colony sampling (top, middle, and base) was done in Florida to determine if *Symbiodinium* genotypes were represented consistently within a colony. Sampling was accomplished using a combination of clipping small pieces of tissue and conducting a biopsy of two or three polyps using the syringe technique (Kemp et al., 2008). All samples were preserved in 95% non-denatured ethanol. Global Positioning System (GPS) coordinates were collected for the Florida colonies so that genetic and clonal diversity estimates could be related to geographic distance among colonies.

**Figure 2.**
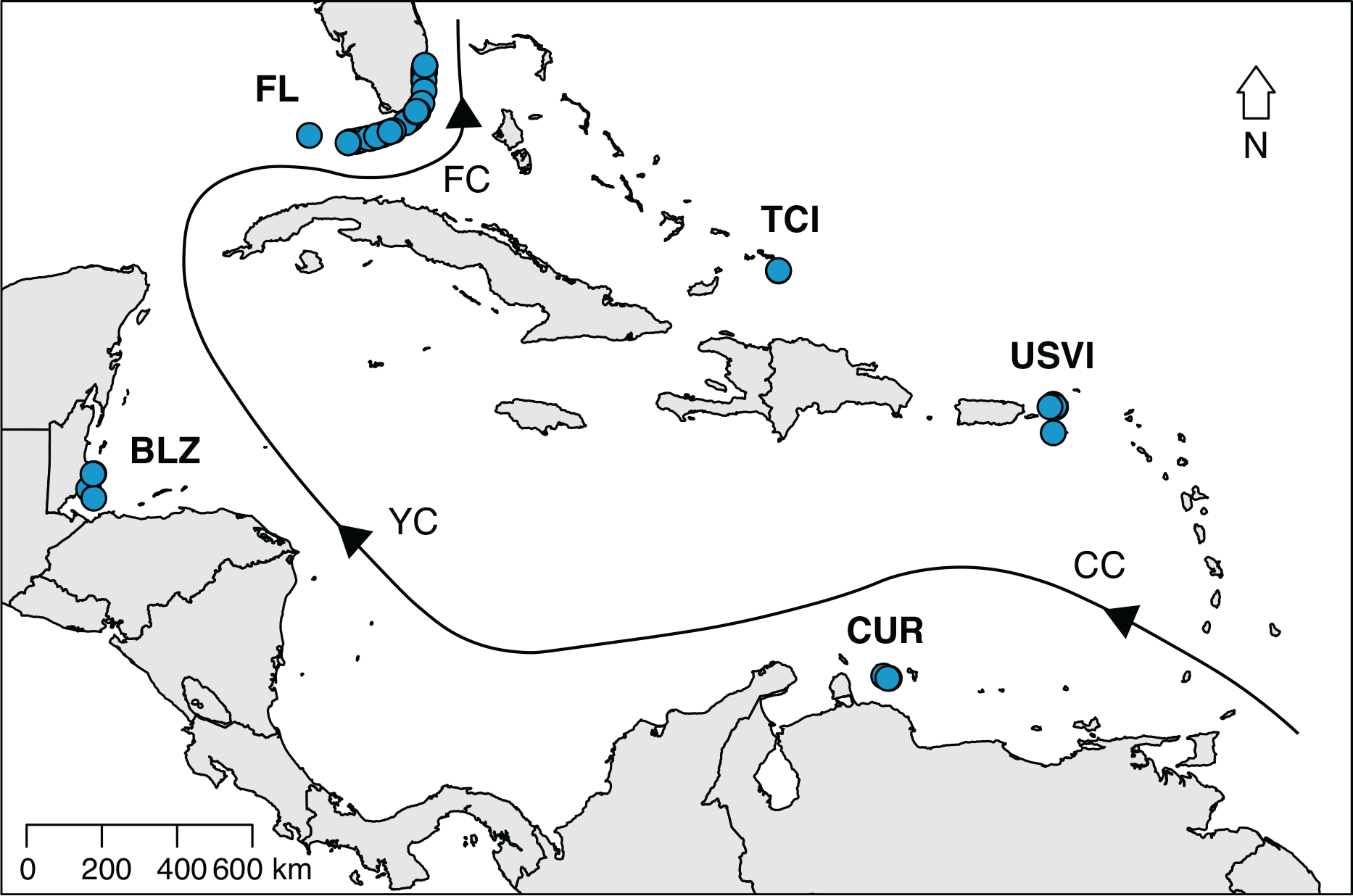
A map of the sites where tissue samples of *Dendrogyra cylindrus* were collected. Samples were collected from 51 sites along the Florida Reef Tract (FL), 3 sites in Curaçao (CUR), 6 sites in the US Virgin Islands (USVI), 3 sites in Belize (BLZ), and 4 sites in the Turks and Caicos Islands (TCI). The black line with arrows indicates the direction of ocean currents. CC=Caribbean Current, YC=Yucatan Current, FC=Florida Current.

### Microsatellite Design and Amplification

One sample from Florida was extracted using the illustra Nucleon PhytoPure Genomic DNA Extraction kit (GE Healthcare) and used as the starting material for generating molecular markers. Because this sample was from an adult colony, DNA from the coral host and dinoflagellate symbiont was sequenced on the Roche GS 454 FLX+ utilizing the Titanium Sequencing Kit (Roche, one half plate). Raw data is available from Penn State’s ScholarSphere (https://scholarsphere.psu.edu/concern/parent/31z40kt37x/file_sets/x6969z327w). See supplement for detail in primer design. After testing primers for successful amplification, host specificity, and variability, 11 markers were retained (Table 1). These markers were combined into four multiplex reactions (labeled A through D, Table 1) using the MultiplexManager software (www.multiplexmanager.com). Only markers labeled with different colored fluorescent dyes (Applied Biosystems) were combined. Multiplex reactions consisted of 1 µl of template DNA, 1.01x Reaction Buffer (New England Biolabs), 2.53 mM MgCl (New England Biolabs), 0.005 mg BSA (New England Biolabs), 0.202 mM dNTPs (Bioline), 0.051 µM of forward and reverse primer (Applied Biosystems), 0.75 U Taq polymerase (New England Biolabs), and nuclease-free water in a total reaction volume of 9.9 µl. Thermocycler parameters included an initial denaturation at 94°C for 5 minutes, 30 cycles of denaturing at 94°C for 20 seconds, annealing at 55°C (or 54°C) for 20 seconds, and extension at 72°C for 30 seconds, and a final extension at 72°C for 30 minutes. All PCR products were visualized using an ABI3730 (Applied Biosystems) automated DNA sequencer with an internal size standard (Gene Scan 500-Liz, Applied Biosystems) for accurate sizing. Electropherograms were analyzed using GeneMapper Software 5.0 (Applied Biosystems).

**Table 1.**
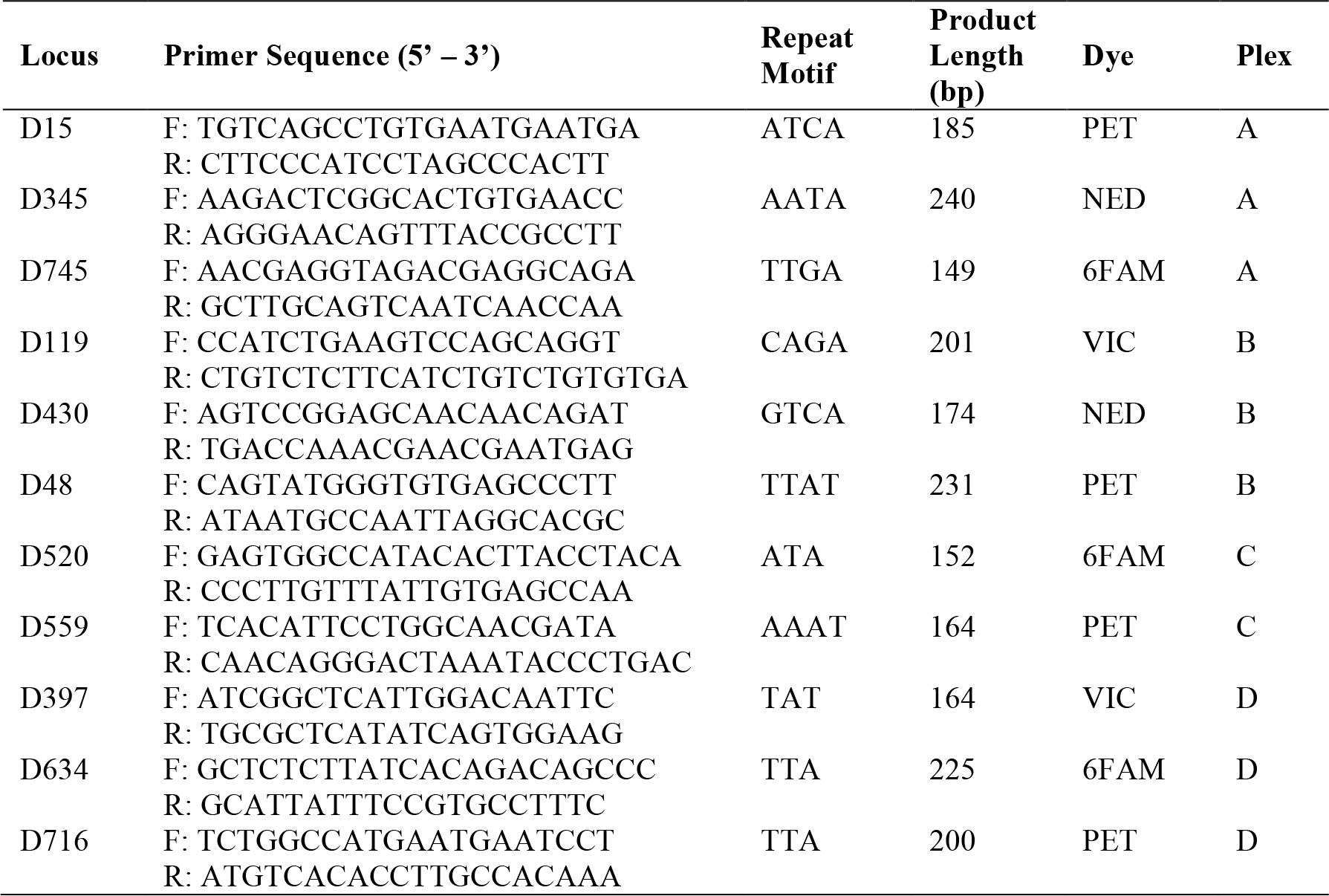
Eleven microsatellite loci developed for *Dendrogyra cylindrus*. Primers with distinct fluorescent dyes were grouped into multiplexes. All reactions were amplified with an annealing temperature of 55°C, except Plex C, which was amplified with a 54°C annealing temperature.

Existing *Symbiodinium* B1 primers (Andras et al., 2009; Pettay & LaJeunesse, 2007; Santos & Coffroth, 2003) were tested on *D. cylindrus* samples from Florida and Curaçao, of which eight were variable and amplified the target product a majority of the time (Table 2). These were combined into multiplex reactions (Table 2, Supplemental Table 1). See the supplement for reaction conditions. Thermocycler parameters varied by multiplex (Supplemental Table 1). However, after conducting preliminary analyses using this set of markers, it became clear that additional markers would be necessary to increase our power of identifying clonal strains. See the supplement for details on *de novo* primer design and amplification conditions. The combined set of 15 microsatellite markers (Table 2) was used to assess clonal and population structure of the algal symbiont.

**Table 2.**
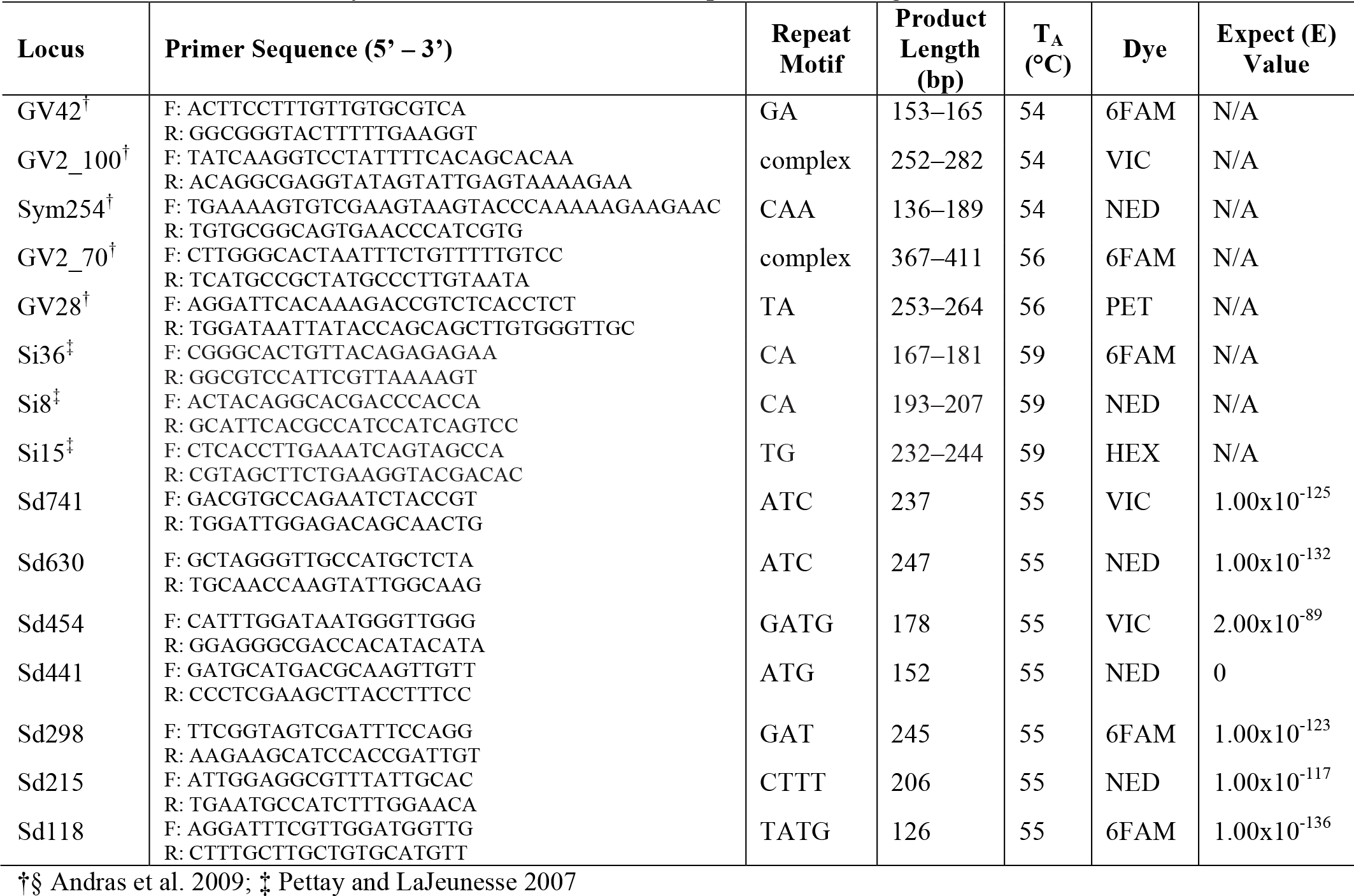
The eight existing and seven *de novo* microsatellite loci used to genotype *Symbiodinium dendrogyrum*. T_A_ = Annealing Temperature. The Expect (E) value indicates the number of matches to the reference that would occur by chance. Low E values correspond to more significant matches.

### Analysis of Population and Clonal Structure

Matching multilocus genotypes were identified using the Data Subset option in GENALEX vers 6.5 (Peakall & Smouse, 2006). Linkage disequilibrium tests were conducted using GENEPOP ON THE WEB 4.0 (Rousset, 2008) Option 2.1. Tests for Hardy-Weinberg equilibrium were conducted in GENALEX vers 6.5 (Peakall & Smouse, 2006) and confirmed using exact tests in GENEPOP ON THE WEB Option 1.1. The probability of identity (PI) describes the power of the microsatellite markers to distinguish closely related colonies (such as siblings) from colonies that are identical because they are the result of asexual reproduction (fragmentation). For the diploid *D. cylindrus*, PI was calculated in GENALEX vers 6.5. The PI for the haploid symbiont was calculated as the sum of each allele’s frequency squared in each population and multiplied across loci.

Genotypic diversity indices for the coral host and symbiont, which included genotypic richness, diversity, and evenness, were calculated as in Baums et al. (2006) using the entire data set of multilocus genotypes (including clones). Unique multilocus genotypes were clustered using the program STRUCTURE v. 2.3.4 to estimate the number of different populations of host and symbiont in the dataset (Pritchard et al., 2000). After initial testing, the admixture model with no location prior was used with correlated allele frequencies. The prior number of populations (*K*) was set from 1 to 6 with five replicate runs per *K*, a burnin of 100,000 and 1,000,000 Markov Chain Monte Carlo repetitions after the burnin using the package ‘PARALLELSTRUCTURE’ (Besnier & Glover, 2013) in R v3.4 (R Development Core Team 2017). The optimal value for *K* was identified using the delta K method (Evanno et al., 2005) implemented in the online program STRUCTURE HARVESTER (Earl, 2012). CLUMPAK was used to identify the consensus of inferred clusters for the different replicates of each *K* value and to visualize the results (Kopelman et al., 2015). Additional estimators of the number of optimal populations (posterior probability, MedMeaK, MaxMeaK, MedMedK, and MaxMedK) were applied (Pritchard et al., 2009; Puechmaille, 2016). Because uneven sampling has been shown to underestimate the true K (Puechmaille, 2016), larger populations were randomly subsampled to create more even sample sizes across the five regions. Global *F_ST_*, or the proportion of genetic variation between subpopulations (Wright, 1951), was calculated using GENODIVE (Meirmans & Van Tienderen, 2004). The *F_ST_* value adjusted for the maximum amount of within-population diversity (*F’_ST_*) was also obtained from GENODIVE. The congruence of population structure and clonal reproduction in *D. cylindrus* and *S. dendrogyrum*, as well as the extent of clonal propagation, were compared.

Analysis of molecular variance (AMOVA) was employed to test hypotheses about population structure of host and symbiont (Excoffier et al., 1992). Samples were grouped by region (identified using STRUCTURE and the optimal *K* value) and location (Belize, Florida, Turks and Caicos Islands, USVI, and Curaçao). The AMOVA option in GENALEX vers 6.5 was used to partition genetic variation from a distance matrix between regions and locations, using 9999 permutations and assuming an Infinite Allele Model.

GPS coordinates for all Florida colonies were used to create a geographic distance matrix, which was compared to the genetic distance matrix for *D. cylindrus*. Mantel tests of isolation by distance and spatial autocorrelation analyses were conducted in GENALEX vers 6.5.

Because of the small number of samples, a finding of no population structure could be due to a lack of power. Thus, simulations were conducted in POWSIM v.4.1 to assess the power of the microsatellite data set to detect low levels of population differentiation. POWSIM estimates the lowest level of differentiation (*F_ST_*) that can be detected in a simulated population with a minimum of 90% accuracy (Ryman & Palm, 2006). Significance was determined using Fisher’s exact test.

### Population Demographic Modeling

The modeling software MIGRAINE v.0.5.2 (http://kimura.univmontp2.fr/~rousset/Migraine.htm) was used to test for past changes in population size in the Florida *D. cylindrus* population. Samples from other locations were not grouped with Florida because population structure within a dataset yields inaccurate demographic parameter inference (Leblois et al., 2014). Additional samples were included for this analysis to increase the power of detecting historical population size changes (n=95 total). The null model assuming a constant population size (OnePop) was run to estimate pGSM, which was close to 0.3. This parameter describes the geometric distribution of mutation step size under a generalized stepwise mutation model (GSM), which is most appropriate for microsatellite markers (Leblois et al., 2014). When we ran the model assuming one past change in population size (OnePopVarSize), we fixed pGSM at 0.3 and inferred the current (2*N*µ) and ancestral (2*N_anc_*µ) population sizes, as well as the duration of the contraction or expansion event (*D*g/2*N*) and the ratio of current to ancestral population size (*N_ratio_*). We assumed a mutation rate (µ) of 5 x 10^-^4 per locus per generation, a value typical for microsatellite loci (Estoup & Angers, 1998). In addition, we also ran the OnePopFounderFlush model, which assumes two past changes in population size and infers a founder population size (2*N_founder_*µ) and two additional population size ratios (*N_cur_N_founder ratio_* and *N_founder_N_anc ratio_*). Significant population size changes were detected using 95% confidence interval estimates of the population size ratios. If the value 1 (indicating the current and ancestral population sizes were identical) was outside of the 95% confidence interval, then the size change was significant. MIGRAINE uses sequential importance sampling algorithms (De Iorio & Griffiths, 2004a, 2004b) to estimate parameters of past demographic changes from population genetic data. We used 2,000 points, 2,000 trees, and four iterations per run for the models with population size changes. The null model was run using 2,000 points, 100 trees, and three iterations. Two independent runs were conducted per model by changing the estimation seed. Additional model settings are included in the supplement (Supplemental Table 2).

### Projected Population Declines

Through the course of this study, we witnessed a severe decline in Florida *D. cylindrus* (Neely et al. in prep). In Broward County, 86% of the known colonies were lost in two years (Kabay, 2016). Severe thermal stress events such as this one are expected to occur annually by 2042 on average under Representative Concentration Pathway (RCP) 8.5 (van Hooidonk et al., 2017). If we extrapolate our data on the number of coral genotypes and symbiont strains in our sample of 145 *D. cylindrus* colonies to the entire Florida population of 610 colonies (Lunz et al., 2016), we would expect to find 210 coral genotypes and 126 symbiont strains. Using our data on the *D. cylindrus* genotype frequencies and *S. dendrogyrum* strain frequencies, we can simulate the decline of Florida *D. cylindrus* with future thermal stress-related disease events. Using the statistical software R, we generated 610 *D. cylindrus* colonies and assigned each colony a coral genotype and symbiont strain based off of the frequency distribution we observed in our sample of 145 colonies. For simplicity, we only assigned one strain per colony whereas in reality some colonies host more than one strain. Thus, the modeled decline in symbiont diversity may be more extreme then what we would actually see in nature.

Because we do not know the actual rate of decline from the most recent thermal stress event, we can simulate three scenarios where 80%, 50%, and 20% of colonies survive each hyperthermal event. This model assumes that there is no sexual reproduction, no establishment of new clonal fragments, and no successful restoration. We also assume that each high temperature event is equally damaging, resulting in the same percent loss of colonies as the previous event.

After running 100 simulations for each of the three rates of decline, we identified the number of thermal stress events that would cause local extinction of *D. cylindrus* in Florida. *D. cylindrus* was considered to be locally extinct once the average number of colonies (across the 100 simulations) remaining after a stress event was below one.

## Results

### Hardy-Weinberg Equilibrium and Linkage Disequilibrium

The 11 *de novo* microsatellite markers developed for *D. cylindrus* yielded few deviations from Hardy-Weinberg equilibrium (Supplemental Table 3). None of the markers significantly deviated from Hardy-Weinberg in more than one location. Few locus pairs showed significant linkage disequilibrium, and none of the significant deviations was found in all locations. Deviations from Hardy-Weinberg equilibrium could not be tested for the symbiont markers, since *S. dendrogyrum* is haploid. Tests for linkage disequilibrium between the *S. dendrogyrum* markers yielded many apparently linked locus pairs. However, this is expected in a species that primarily reproduces asexually within its host, and thus recombination may be rare. Most paired combinations of the symbiont loci were in equilibrium in at least one location, indicating that these markers are not physically linked. Other locus pairs showed significant linkage disequilibrium in just one location, without enough information available in the other locations to discount linkage equilibrium.

### Genetic Diversity of *de novo Dendrogyra cylindrus* microsatellite markers

The microsatellite markers developed for *Dendrogyra cylindrus* ranged in allelic diversity from 6 to 15 and in effective allelic diversity from 2.047 to 6.849 (Table 3). Observed heterozygosity levels ranged from 0.399 to 0.954, expected heterozygosity within subpopulations ranged from 0.533 to 0.888, and total heterozygosity ranged from 0.686 to 0.895 (Table 3). The total heterozygosity adjusted for sampling a limited number of populations ranged from 0.724 to 0.897 (Table 3). The heterozygosity values for all markers were relatively high (closer to the maximum value of 1), indicating high genetic variability for these markers (Table 3). Inbreeding coefficients for the 11 markers were between −0.19 and 0.361, with most values close to zero (Table 3). The new markers differed in their genetic variability, with D15 showing the lowest differentiation across all four measures (Table 3). D716 and D48 showed the most variability. Overall, the markers were polymorphic enough to assess clonal and population structure in *D. cylindrus*.

**Table 3.** Summary statistics per locus for 11 *de novo* microsatellite markers for *Dendrogyra cylindrus*. *N*_a_=number of alleles; *N*_eff_=effective number of alleles; *H*_o_=observed heterozygosity; *H*_s_=heterozygosity within populations; *H*_t_=total heterozygosity; *H*’_t_=corrected total heterozygosity; *G*_is_=inbreeding coefficient; *G*_st_=fixation index; *G*’_st_(Nei)=Nei, corrected fixation index; *G*’_st_(Hed)=Hedrick, standardized fixation index; *D*_est_=Jost’s D differentiation. Standard errors were calculated by jackknifing over loci. All metrics were calculated using Genodive (Meirmans & Van Tienderen, 2004).

| Locus | $N_a$ | $N_{eff}$ | $H_o$ | $H_s$ | $H_t$ | $H'_t$ | $G_{is}$ | $G_{st}$ | $G'_{st}(\text{Nei})$ | $G'_{st}(\text{Hed})$ | $D_{est}$ |
| --- | --- | --- | --- | --- | --- | --- | --- | --- | --- | --- | --- |
| D15 | 15 | 6.849 | 0.891 | 0.888 | 0.895 | 0.897 | -0.004 | 0.008 | 0.010 | 0.091 | 0.083 |
| D345 | 13 | 3.704 | 0.772 | 0.759 | 0.815 | 0.829 | -0.017 | 0.069 | 0.085 | 0.339 | 0.290 |
| D745 | 10 | 3.375 | 0.749 | 0.731 | 0.789 | 0.803 | -0.025 | 0.073 | 0.090 | 0.322 | 0.268 |
| D119 | 7 | 2.977 | 0.611 | 0.694 | 0.749 | 0.763 | 0.119 | 0.074 | 0.091 | 0.284 | 0.227 |
| D430 | 9 | 3.421 | 0.721 | 0.736 | 0.761 | 0.767 | 0.021 | 0.032 | 0.040 | 0.145 | 0.116 |
| D48 | 9 | 2.454 | 0.399 | 0.625 | 0.787 | 0.828 | 0.361 | 0.206 | 0.245 | 0.636 | 0.541 |
| D520 | 13 | 5.861 | 0.865 | 0.862 | 0.882 | 0.887 | -0.004 | 0.022 | 0.028 | 0.197 | 0.178 |
| D559 | 6 | 3.036 | 0.732 | 0.696 | 0.758 | 0.773 | -0.052 | 0.082 | 0.100 | 0.316 | 0.255 |
| D397 | 11 | 3.865 | 0.690 | 0.774 | 0.795 | 0.800 | 0.109 | 0.027 | 0.033 | 0.140 | 0.117 |
| D634 | 11 | 4.486 | 0.954 | 0.802 | 0.815 | 0.818 | -0.190 | 0.016 | 0.020 | 0.098 | 0.083 |
| D716 | 6 | 2.047 | 0.506 | 0.533 | 0.686 | 0.724 | 0.051 | 0.223 | 0.264 | 0.541 | 0.409 |
| Overall | 10.00 | 3.825 | 0.717 | 0.736 | 0.794 | 0.808 | 0.026 | 0.072 | 0.089 | 0.325 | 0.273 |
| SE | 0.894 | 0.432 | 0.050 | 0.030 | 0.018 | 0.016 | 0.038 | 0.021 | 0.026 | 0.064 | 0.052 |

$G_{st}$ is a measure of genetic differentiation among populations that is generalized for markers with multiple alleles, and is analogous to $F_{st}$ . $G'_{st}$ The adjustment to this statistic formulated by Nei corrects for sampling a small number of populations (Nei, 1987). $G'_{st}(\text{Hed})$ Hedrick's $G'_{st}$ is standardized relative to the maximum level of differentiation based on the heterozygosity within subpopulations (Hedrick, 2005). $D_{est}$ Contrastingly, Jost's measure of population differentiation is independent of the within subpopulation diversity (Jost, 2008).

### Clonal Structure and Spatial Analyses along the Florida Reef Tract

Clonal structure analyses for the coral host revealed that the sampled colonies of *D. cylindrus* along the Florida Reef Tract were highly clonal within each site. After genotyping 145 colonies from 47 different sites, we only found 50 unique multilocus genotypes (Figure 3). The Florida population of the coral host also had low values for genotypic richness, diversity, and evenness (Figure 3). This indicated that within each site, the Florida coral colonies were predominantly the product of asexual reproduction. The host probability of identity values for all sampling regions were less than 1.0×10^−10^, indicating a low probability of misidentifying clones. The probability of identity values for the symbiont were all reasonably low (Waits et al., 2001), with the exception of the Turks and Caicos Islands region (Curaçao: 0.0036, USVI: 0.015, Florida: 1.5×10^−5^, Belize: 0.003, Turks and Caicos: 0.092). Because sampling was biased towards increasing the number of unique genotypes in all regions except Florida, clonal structure could not be described elsewhere.

**Figure 3.**
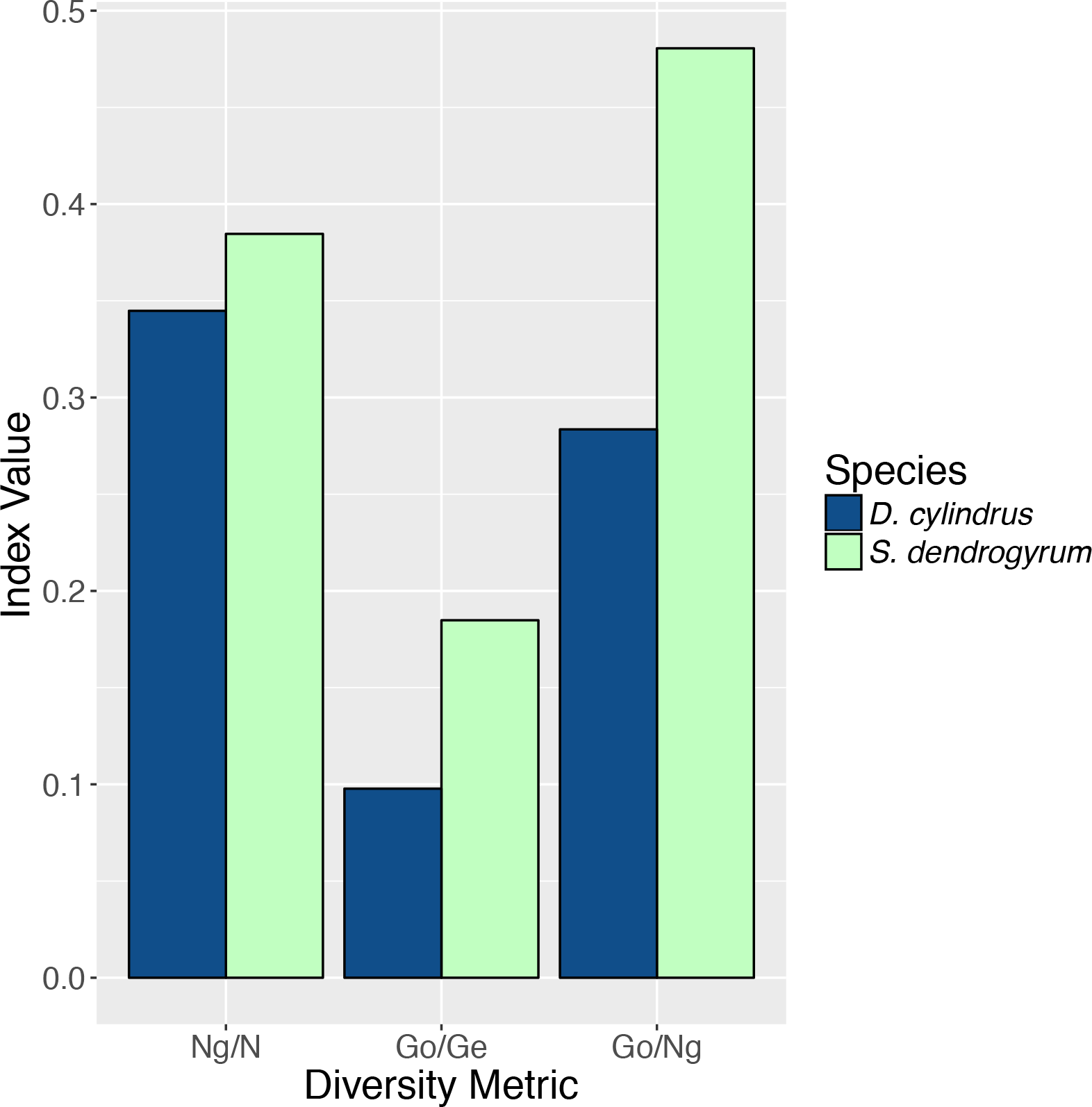
Summary of the clonal structure of *Dendrogyra cylindrus* and its dominant algal symbiont *Symbiodinium dendrogyrum* in Florida. Genotypic richness (Ng/N), diversity (Go/Ge), and evenness (Go/Ng) are plotted on the x-axis and the value of the diversity index is plotted on the y-axis. Ng = number of genotypes, N = number of colonies sampled, G_O_ = observed genotypic diversity, G_E_ = expected genotypic diversity.

*S. dendrogyrum* is haploid, and thus all samples with more than one allele were considered to be multiple infections. There was no spatial pattern in samples with single versus multiple infections along the Florida Reef Tract (Supplemental Figure 1). These samples were not included in further analyses. After removing samples with multiple infections (n=98) and samples that failed in one or more of the 15 microsatellite markers (n=97), we obtained a subset of samples (n=112) with complete multi-locus genotypes. Only 30 unique genotypes were found in Florida, with most strains confined to single reefs. The values for genotypic richness standardized to sample size (Ng/N), genotypic diversity (Go/Ge), and genotypic evenness (Go/Ng) were all higher in the algal symbiont, compared to the host (Figure 3). The richness values were the most similar, indicating that we found a similar number of unique genotypes of host and symbiont relative to the number of samples we genotyped for each species. The values for genotypic diversity were low in both host and symbiont (albeit, slightly higher in the symbiont), demonstrating that both species reproduced mostly asexually within a site. Genotypic evenness was higher in the symbiont relative to the host, meaning that the host population was dominated by relatively few highly replicated genotypes.

For the coral host, all of the clonal ramets of the same genet were contained within a single collection site. This indicates limited dispersal abilities of asexual fragments of *D. cylindrus*. A single coral genet dominated most of our sites, however, three sites had two genets and one site contained three different genets (Figure 4A).

**Figure 4.**
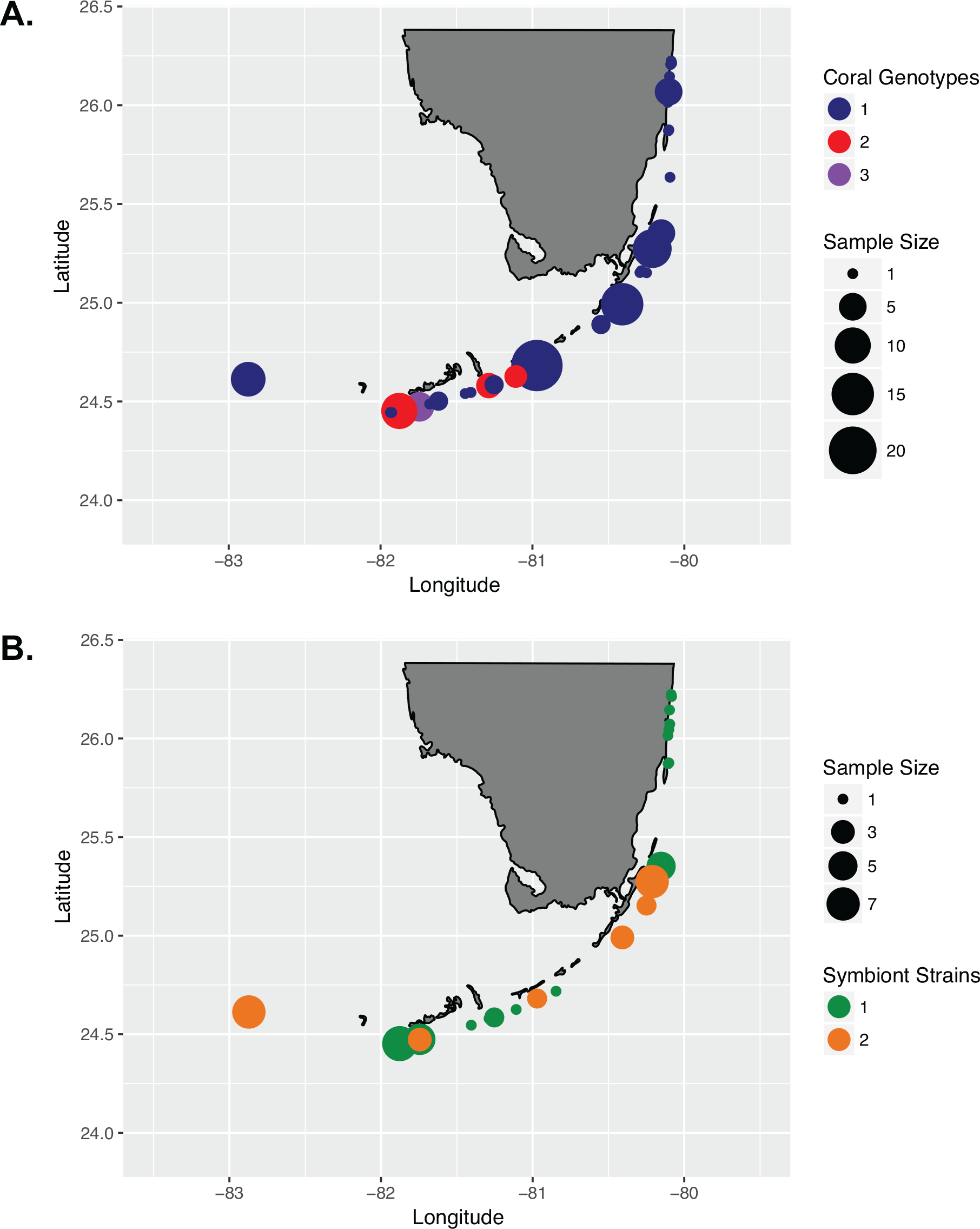
Maps of Florida collection sites of *Dendrogyra cylindrus* (A) and *Symbiodinium dendrogyrum* (B), showing sites with one coral genet (blue), two coral genets (red), and three coral genets (purple). Sites with one symbiont strain are shown in green, and sites with two symbiont strains are shown in orange. The size of the circle corresponds to the relative number of samples genotyped from each site. Note the differing scales.

A test of isolation by distance for *D. cylindrus* along the Florida Reef Tract (n=50) revealed no significant correlation between genetic and geographic distance (r^2^ = 0.0001). Spatial autocorrelation analysis using the complete dataset of *D. cylindrus* genotypes (including clones) in Florida (n=180) revealed significant positive spatial autocorrelation up to distances of 60 meters (Supplemental Figure 2A). However, when clones were removed, there was no significant spatial autocorrelation (Supplemental Figure 2B). These results indicate that the cause of the correlation between genetic distance and geographic distance over small distance classes is a result of asexual reproduction, likely via fragmentation, in *D. cylindrus*.

The genotype of the algal symbiont often corresponded with the genotype of the coral host, meaning that all of the ramets of a coral host genet were often symbiotic with the same clonal strain of *S. dendrogyrum*. There were exceptions to this observation. For example, in the Dry Tortugas, all of the colonies were ramets of the same coral host genet. However, different ramets of this genet associated with two distinct strains of *S. dendrogyrum* (Figure 4). The reverse was also observed, with the same symbiont strain found within different coral genets at different sites. The same symbiont strain was found at both Bahia 3 and Bahia 4, which are 183 meters apart and contain corals belonging to different genets. One coral colony at Marker 32-3 had the same symbiont strain as the coral colonies at Marker 32-1 (50 meters apart), despite there being different coral genets at these sites. The different coral genets at Hollywood and PC Ledge contained the same symbiont strain as well (3.44 km apart). The same was true for Pompano 1 and Pompano Drop-off (1.23 km apart). All of these pairs of sites that share identical symbiont strains are relatively close together (Figure 5). We also sampled the same coral colony in different locations (e.g. top, middle, and base) to assess symbiont diversity. Of the five colonies that were sampled in this way and successfully genotyped, only one contained two different symbiont strains.

**Figure 5.**
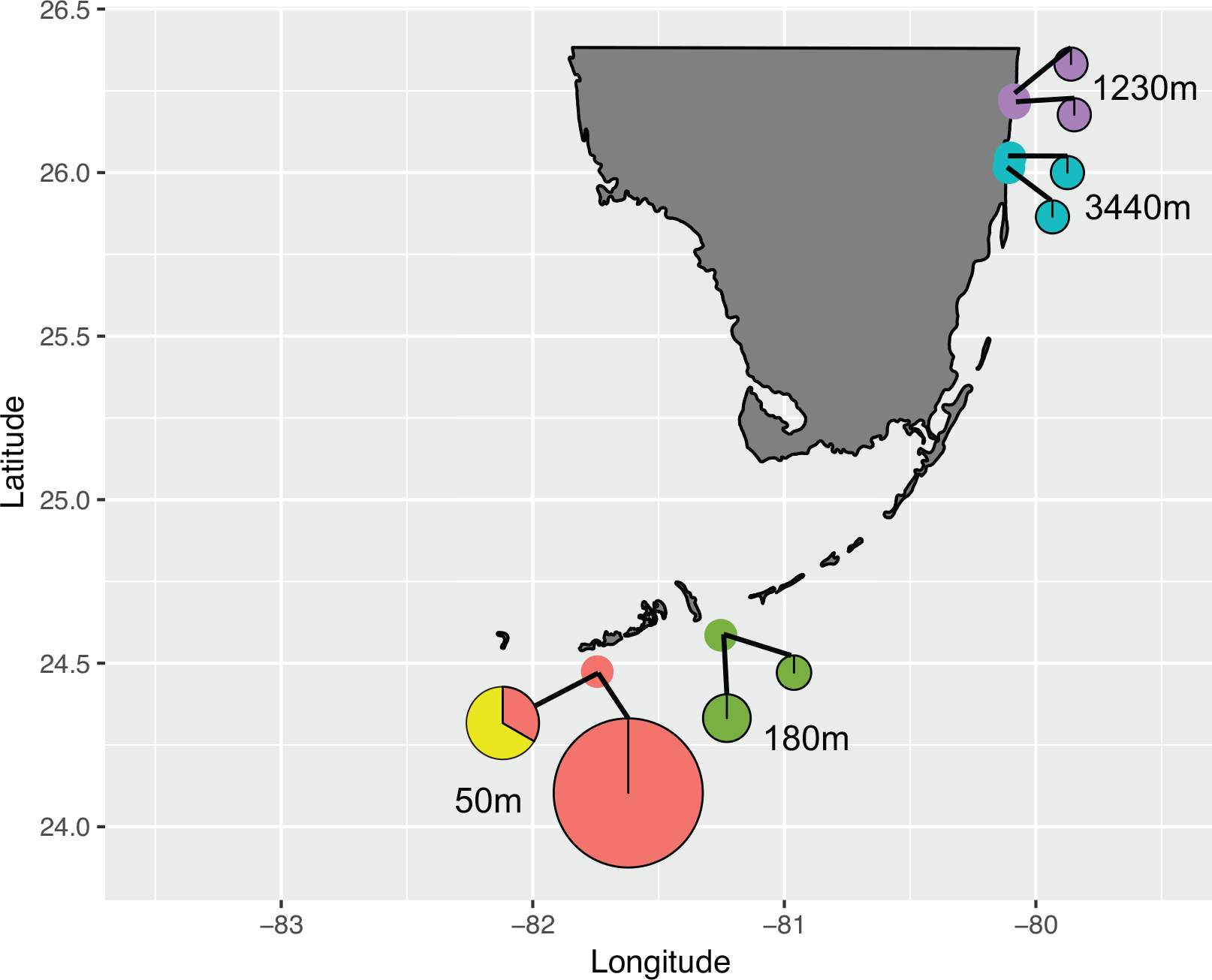
Four pairs of sites that shared identical symbiont strains are shown along the Florida Reef Tract. Each pie chart corresponds to a site, with a line connecting the chart to the site’s geographic location represented by a point with the same color. The colors of the pie chart sections correspond to the strain composition at that site, with color indicating symbiont identity. The size of the pie charts corresponds to the number of symbiont samples genotyped from that site. The distance between each pair of sites is indicated on the map in meters.

### Population Structure Analyses

Strong genetic differentiation between populations was found, with a significant global *F_ST_* value of 0.110 and an *F’_ST_* value of 0.414. Bayesian clustering using the entire dataset of unique multilocus genotypes for the coral host revealed the presence of two to four clusters in the dataset. The different estimators for the optimum number of populations yielded different results. For example, the ΔK method (Evanno et al., 2005) yielded a K of 2 and the MaxMeaK with a spurious cluster threshold of 0.5 (a less conservative measurement) yielded a K of 4. In the STRUCTURE plot for K=2, all of the Florida, Belize, and Turks and Caicos samples have a high probability of membership to the first cluster and all of the Curaçao and US Virgin Islands samples have a high probability of membership to the second cluster (Figure 6A). When the same dataset was run in STRUCTURE setting K=3 as the *a priori* number of populations in the dataset, an additional cluster corresponding to the individuals from Curaçao separates from the US Virgin Islands (Figure 6B). The samples from the Turks and Caicos Islands also appear to be admixed. However, our dataset contained locations that were sampled unevenly. Thus, we randomly selected unique multilocus genotypes to a maximum size of 20 to remove the bias associated with uneven sampling (Puechmaille, 2016). When Bayesian clustering analysis was completed for this subsampled dataset, all of the K estimators yielded an optimal cluster number of three. The only exception was the ΔK method, which still indicated that there were two populations. An AMOVA corroborated these results, with the largest pairwise *F_ST_* value between Florida and Curaçao (*F_ST_*=0.184, p-value<0.001) indicating that these populations of *D. cylindrus* have been separated for a long time. The US Virgin Islands and Curaçao were significantly differentiated, although less strongly (*F_ST_*=0.045, p-value<0.001). The Turks and Caicos Islands were equally differentiated from Florida and the US Virgin Islands (*F_ST_*=0.046, p-value<0.001), further supporting that this is likely an area of admixture.

**Figure 6.**
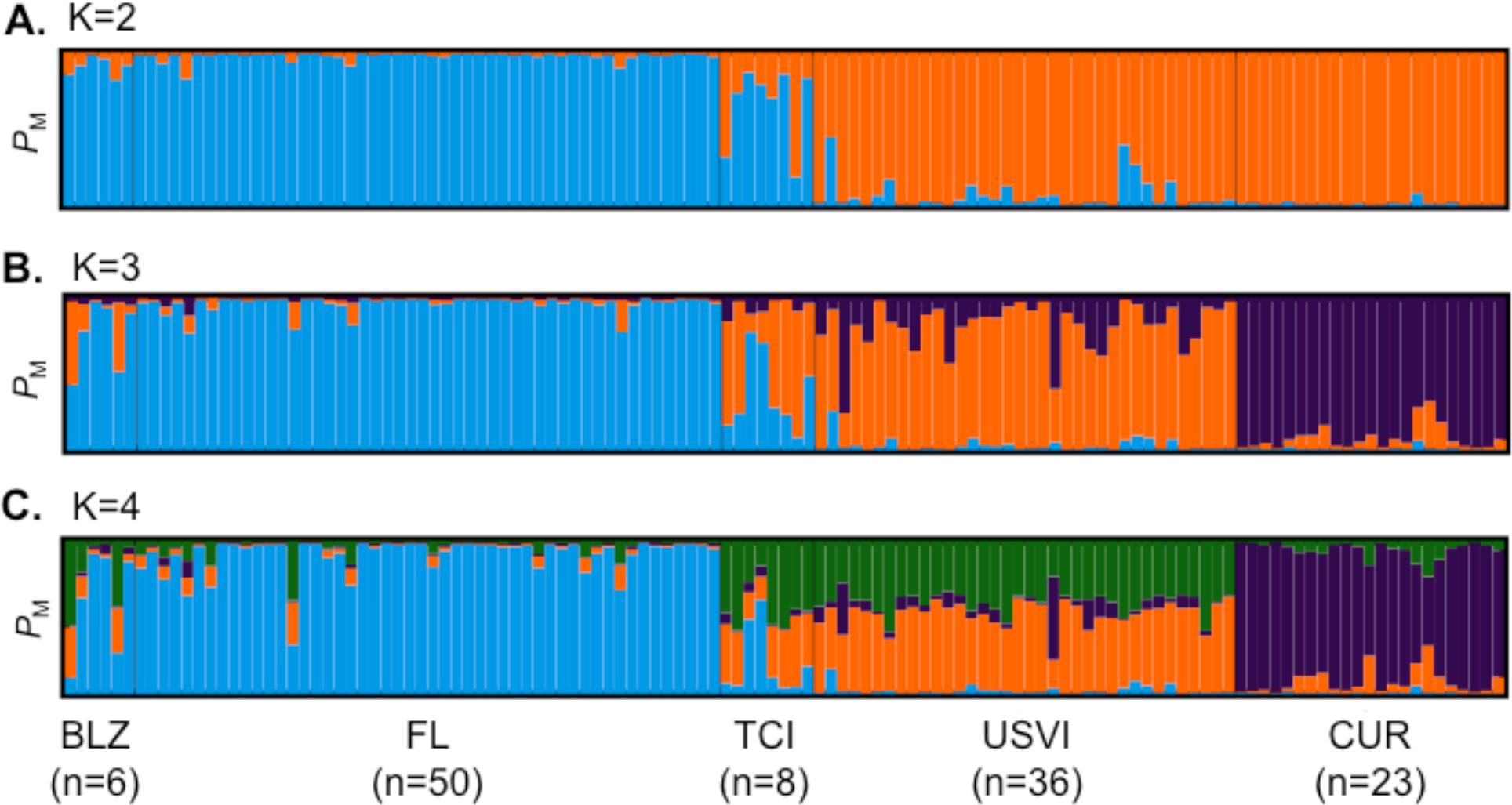
Population structure results for *Dendrogyra cylindrus*. Each vertical bar represents a unique multilocus genotype. Probability of membership to a cluster is plotted on the y-axis. **A**. STRUCTURE results for *D. cylindrus* revealed two separate clusters (K=2, indicated by color). **B.** STRUCTURE results for *D. cylindrus* when K=3. **C.** STRUCTURE results for *D. cylindrus* when K=4. Individuals are grouped by sample location; abbreviations are the same as in Figure 2.

We did not find significant population structure within Florida for the coral host. POWSIM v.4.1 with 1000 simulation runs using the allele frequencies from our host marker set was used to test whether this result was due to a lack of power. Fisher’s exact test revealed a high statistical power (1-β=0.97) of detecting an *F_ST_* value of 0.0195, and a low type I error rate (α=0.0410). Thus, our sample size of 50 individuals from Florida was adequate to detect low levels of population structure along the FRT if present.

Population structure analyses were also conducted for *S. dendrogyrum*. Only 58 unique genotypes were found, after removing samples with multiple infections and clones. Cluster analysis using STRUCTURE and a consideration of multiple estimators for the optimum number of populations revealed that the likely number of populations in the dataset is between 2 and 4 (Figure 7). However, there were locations in this dataset that were sampled more intensely than others, and thus we subsampled the locations with the largest sample size to a maximum value of 19. Because we found population structure for *S. dendrogyrum* within Florida, we subsampled by the five regions (i.e. Broward, Upper Keys, Middle Keys, Lower Keys, and Dry Tortugas) and included the only sample from Biscayne. Most of the K estimators indicated that there were likely four populations in the dataset. When the threshold for spurious clusters was set to a strict 0.8, some of the K estimators yielded a K of 2. The ΔK method alone indicated that the optimal number of populations was three. An AMOVA for the haploid *S. dendrogyrum* revealed that the highest amount of between-group variation was between sample locations (*ϕ_PT_*=0.520, p-value=0.001).

**Figure 7.**
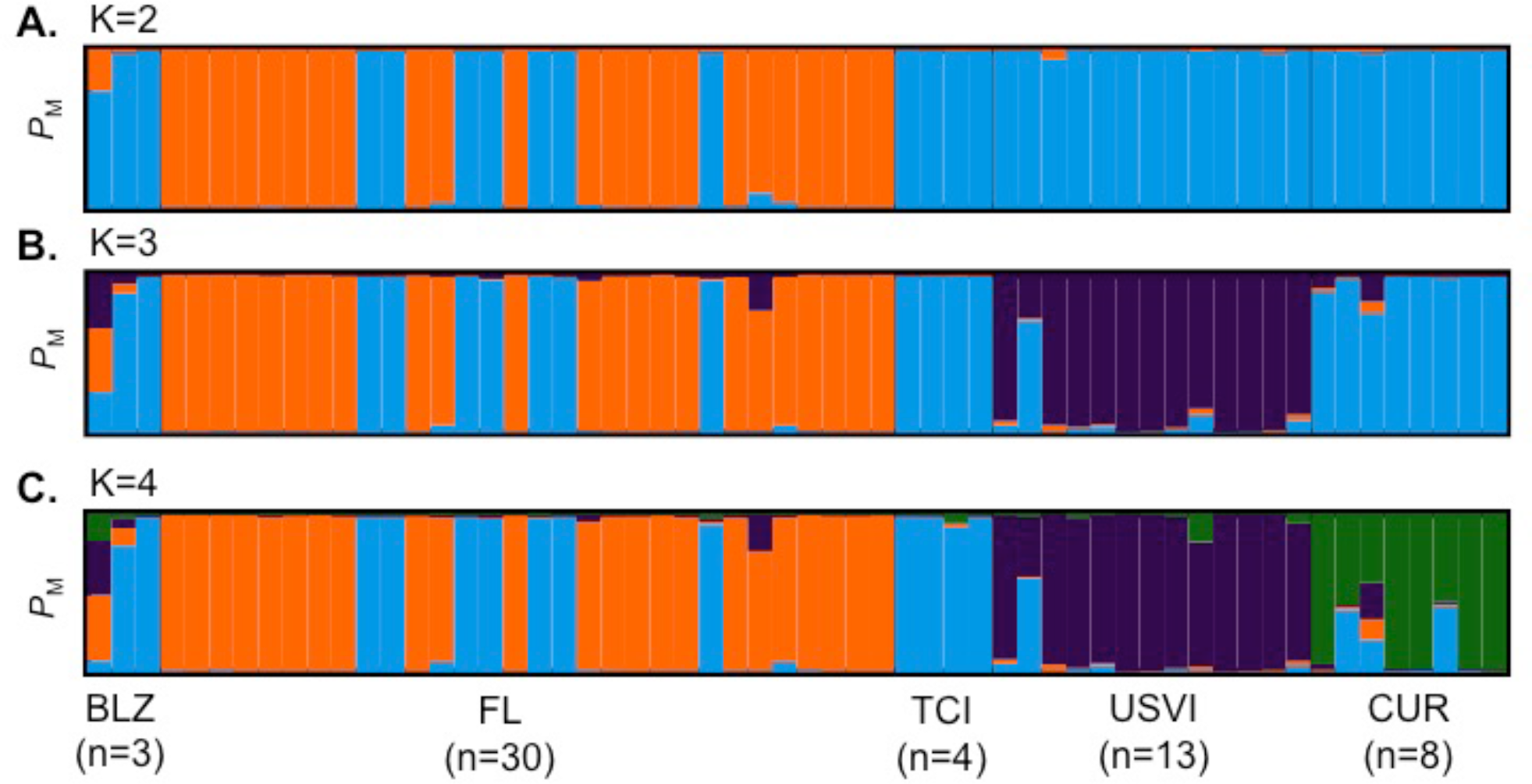
Population structure results for *Symbiodinium dendrogyrum*. Each vertical bar represents a unique multilocus genotype. Probability of membership to a cluster is plotted on the y-axis. **A**. STRUCTURE results for *S. dendrogyrum* when K=2. **B.** STRUCTURE results when K was set to 3. **C.** STRUCTURE results when K was set to 4. Individuals are grouped by sample location; abbreviations are the same as in Figure 2.

These results indicate that the population structure of the algal symbiont did not match the population structure of the host. Notably, there was population structure within the FRT for the symbiont that was not found in the host. Some samples from Florida clustered more strongly with samples from Belize and the Turks and Caicos Islands than other Florida samples. Because this pattern of population assignment did not correspond with geography, four samples from the unique Florida cluster and three samples from Florida that clustered with the rest of the Caribbean were sequenced using microsatellite flanker sequences (Si15) to confirm that they all belonged to the same species. Comparing the Si15 sequences to representative sequences from *Symbiodinium* Clade B (Finney et al., 2010) revealed that samples from the two clusters within Florida are indeed the same species (*S. dendrogyrum*). Because such a small sample of symbiont strains was included from both Belize and the Turks and Caicos Islands, it is possible that the cluster containing these samples and some Florida samples is an artifact of STRUCTURE (Puechmaille, 2016).

### Historical Changes in Pillar Coral Population Sizes

Demographic modeling results revealed no evidence for past changes in pillar coral population size. In both the one continuous population size change (OnePopVarSize) and the two population size changes (OnePopFounderFlush) models, the upper limit of the 95% confidence intervals for the ancestral population size (2*N_anc_*µ, both models) and the founder population size (2*N_founder_*µ, OnePopFounderFlush model only) were undefined. The undefined upper limit meant that many possible population sizes for the ancestral and founder population were equally likely, including population sizes that were both larger and smaller than the 95% confidence interval for the current population size (2*N*µ). This was further evidenced by the broad profile likelihood ratios for the ancestral and founder population sizes (Supplemental Figure 3), and the population size ratios that included the value 1, indicating no significant change. To decide which of the three models best described our data, we used the Akaike Information Criterion (AIC) calculated based on the log likelihood and the number of parameters in each model (Anderson, 2007). The AIC values for the OnePop (798.94), OnePopVarSize (799.56), and OnePopFounderFlush (800.6) models were very close to one another. Because lower AIC values indicate better model fit, we conclude that the OnePop model with no past change in population size is the best model. However, the differences in AIC between models were very small, and thus it is possible that all three models do not accurately describe reality. Still, the results from the models assuming past demographic events failed to detect significant changes in population size, thus supporting our conclusion of no past changes in *D. cylindrus* populations.

### Projected Population Declines

The three projected rates of decline yielded a range of times to extinction for *D. cylindrus*. In the most conservative simulated scenario, where 80% of colonies survived each hyperthermal event, it took 29 stress events for extirpation of *D. cylindrus* from the FRT. With 50% and 20% survival, it took ten and four stress events, respectively, for local extinction to occur. The results from simulations assuming 50% survival are shown in Figure 8. Because some coral colonies are ramets of the same genet, and these coral ramets often contain clonal strains of *Symbiodinium*, coral colonies are lost at a faster rate than coral genotypes and symbiont strains (Figure 8).

**Figure 8.**
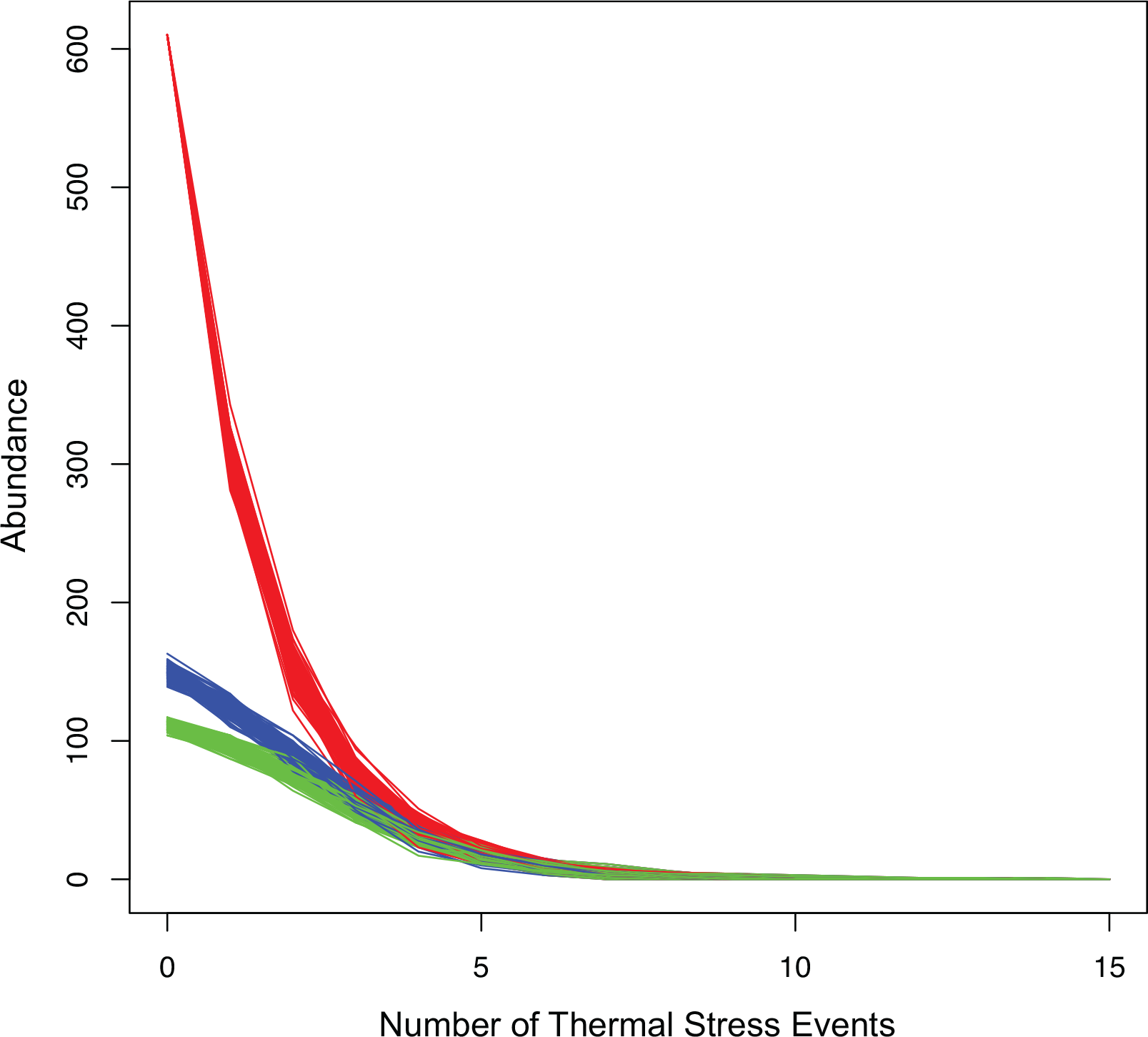
Results from simulating the projected decline of *Dendrogyra cylindrus* in Florida, assuming a survival rate of 50% with each severe thermal stress-related disease outbreak. The red lines correspond to the number of *D. cylindrus* colonies, the blue lines indicate the number of unique coral genotypes, and the green lines show the decline in the number of unique strains of *Symbiodinium dendrogyrum*. Each line is the outcome of one simulation, for a total of 100 simulations.

## Discussion

The pillar coral, *Dendrogyra. cylindrus* and its obligate algal symbiont *Symbiodinium dendrogyrum* reproduce locally via asexual processes albeit dispersal distances of their asexual propagules differ, with strains of the algae dispersing over somewhat larger scales. Barriers to gene flow were also not congruent between the partners but showed similarities with other coral-algal symbioses in the Caribbean. As is often the case for corals (Baums, 2008), while asexual reproduction was prevalent, no signs of inbreeding were detected in the coral host and allelic diversity was relatively high. Demographic modeling based on molecular data agreed with the geological record that *D. cylindrus* was historically not a dominant species on Caribbean reefs and yet was able to survive as the sole remaining species in the genus. The forecasted increasing frequency of extreme warm water events in combination with the absence of sexual recruitment in this species project a high likelihood that this species will be the first coral to become locally extinct in the Florida Keys in modern times. The consequences of the loss of rare coral species and their symbionts are unknown but evidence from other ecosystems indicates that loss of rare species can destabilize communities and degrade ecosystem function.

### Clonal Structure

Over short distances (<60 m), Florida *Dendrogyra cylindrus* colonies were highly clonal and showed positive spatial autocorrelation, indicating that the primary mode of reproduction over this scale is asexual fragmentation (Supplemental Fig 2, Figure 3). These clonal structure results are similar to those of the elkhorn coral, *Acropora palmata* (Baums et al., 2006) and the massive coral, *Orbicella faveolata* (M. Miller et al., 2018). The prevalence of asexual reproduction in Florida *A. palmata* was attributed to increased retention of fragments on the larger shelf area, and low rates of sexual recruitment. Similar forces are likely responsible for high asexual reproduction in Florida *D. cylindrus*.

In benthic surveys of nine inshore to offshore sites in Key Largo, Florida, no juvenile colonies of *D. cylindrus* were discovered (S. L. Miller et al., 2010), indicating successful sexual recruitment of juvenile pillar corals is nonexistent or very low. In addition, the Florida populations of both *D. cylindrus* and *A. palmata* are at the northern edge of these species’ ranges. Previous work on clonal plants and corals has pointed to the prominence of asexual reproduction at the edge of species’ geographical ranges, enabling marginal populations to persist despite low sexual recruitment (Boulay et al., 2014; Silvertown, 2008). Thus, recent reproduction in the Florida *D. cylindrus* population may be entirely asexual due to the lack of sexual recruits from neighboring populations and from within Florida.

Because most sites tended to be occupied by a single coral genotype (Figure 4A), it is possible that Florida *D. cylindrus* are already experiencing an Allee effect, wherein the density of compatible colonies is too low for successful sexual reproduction and population growth (Courchamp et al., 1999). This study was limited to assess clonal structure in Florida alone, and thus future work should measure genotypic richness, diversity, and evenness in other locations.

The Florida population of *S. dendrogyrum* is also clonal, with the same symbiont strain often (but not always) found in all of the ramets of a coral genet. High fidelity between individual genotypes of symbiotic partners has been found previously in the Caribbean elkhorn coral (*Acropora palmata*) and its algal symbiont (*Symbiodinium ‘fitti’*) (Baums et al., 2014b). While coral genets were restricted to one site each, five pairs of sites did share symbiont strains. The sites that shared symbiont strains were close together (Figure 5). Thus, the symbiont may be able to disperse asexually over larger distances than its coral host, although successful long-distance dispersal appears to be rare. A typical fragment dispersal distance for *D. cylindrus* is less than 60 meters, as evidenced by the spatial autocorrelation analysis (see below and Supplemental Figure 2).

Similarly to the situation in *D. cylindrus* and *S. dendrogyrum*, the asexual dispersal ability of *Symbiodinium fitti* was found to be significantly higher than its coral host, *A. palmata* (Baums et al., 2014b). However, the actual asexual dispersal distances for *S. dendrogyrum* were lower than what was found for *S. fitti* (greater than 2000 meters). While *Symbiodinium* cells are constantly being expelled from the coral host, little is known about their free-living stage. Despite similar morphologies, different species of *Symbiodinium* may differ greatly in their abilities to disperse and infect new host corals.

Populations of foundation species with low genotypic diversity are more susceptible to disturbances. Increasing genotypic diversity in plots of the seagrass, *Zostera marina*, yielded higher resistance to disturbance by grazing geese (A. R. Hughes & Stachowicz, 2004). High intraspecies variation in crop plants increases plant fitness and agricultural yields, and decreases susceptibility to insect pests (Tooker & Frank, 2012). In addition, high genotypic diversity confers a greater ability for populations to recover from warm temperature stress events (Reusch et al., 2005), which are expected to become more frequent with climate change (Stocker et al., 2013).

### Barriers to Gene Flow

Population structure results for *D. cylindrus* indicate that there are two genetic breaks in the Caribbean, resulting in three populations. The first population includes samples from Florida, Belize, and the Turks and Caicos Islands, the second population includes samples from the U.S. Virgin Islands, and the third population comprises samples from Curaçao (Figure 6). Cluster analysis results and pairwise *F*_ST_ analyses revealed that there is little to no gene flow between Florida and either Curaçao or the USVI. A genetic break in this area has been well characterized in other corals (Andras et al., 2013; Baums et al., 2010; Foster et al., 2012; Rippe et al., 2017; Vollmer & Palumbi, 2006).

*Dendrogyra cylindrus* lacks population structure along the Florida Reef Tract, which was also found in both *A. cervicornis* and *A. palmata* using microsatellite markers (Baums et al., 2010; Baums et al., 2005). However, an analysis of Florida *A. cervicornis* using SNPs found significant population structure within the Florida Reef Tract (Willing et al., 2012), thus it is possible that higher resolution markers may resolve additional structure in *D. cylindrus*.

The genetic break between *D. cylindrus* in the USVI and Curaçao was weaker than the break between Florida and these two locations but again consistent with a similar genetic break in *A. palmata* (Devlin-Durante & Baums, 2017). It is possible that the Caribbean Current is restricting coral larval exchange between these two island nations (Figure 2). This current was shown to affect dispersal in other planktonic marine species (Díaz-Ferguson et al., 2010; Jossart et al., 2017), however, population genetic data for other coral species does not show a break in this area (Andras et al., 2013; Rippe et al., 2017).

After correcting for uneven sampling effort, the results for *S. dendrogyrum* population structure revealed four populations in the Caribbean (Figure 7). *S. dendrogyrum* strains in both Curaçao and the US Virgin Islands separate as two populations distinct from the rest of the Caribbean. These genetic breaks are in concordance with the breaks observed in the host coral, *D. cylindrus*. However, there is population structure within the FRT for the algal symbiont, with some Florida individuals clustering with Belize and the Turks and Caicos Islands. A similar incongruence between the population structure of host and symbiont was found in *G. ventalina* (Andras et al., 2011; Andras et al., 2013) and *A. palmata* (Baums et al., 2014b; Baums et al., 2005). A recent review explained the observed lower connectivity in horizontally-transmitted algal symbionts relative to their coral hosts as resulting from restricted dispersal and recruitment (Thornhill et al., 2017). The inconsistency in population structure between host and symbiont supports that *D. cylindrus* obtains its symbionts horizontally (from the surrounding environment) and implies that gene flow occurs over different spatial scales in the partners (Baums et al., 2014b).

### Demographic Modeling

Demographic modeling results using MIGRAINE revealed no evidence for past changes in population size in Florida *D. cylindrus*. This is consistent with previous descriptions of *D. cylindrus* as a naturally rare species (NOAA, 2014), and with the rarity of *D. cylindrus* in the fossil record (see Introduction). Abundance can be high in localized areas, however, due to asexual fragmentation. This suggests that historically, *D. cylindrus* did not experience a range-wide decline since its first appearance in the fossil record during the late Pliocene/early Pleistocene (Budd, 2000).

Through the course of this study, we witnessed a severe decline in Florida *D. cylindrus* (Neely et al. in prep). During the summers of 2014 and 2015, water temperatures rose above30.5°C in the Florida Keys (http://coralreefwatch.noaa.gov), triggering a multi-year thermal stress event (Lewis et al., 2017; Manzello, 2015). After some coral colonies bleached in 2014, many contracted white plague disease (Precht et al., 2016). This thermal stress-related disease outbreak led to a massive loss of Florida pillar corals (Neely et al. in prep). Some areas experienced higher mortality rates than others. In Broward County, 86% of the known colonies were lost in two years (Kabay, 2016). Severe hyperthermal events such as this one are expected to occur annually by 2042 on average under Representative Concentration Pathway (RCP) 8.5 (van Hooidonk et al., 2017), although there is projected heterogeneity along the Florida Reef Tract (van Hooidonk et al., 2015).

Prior to the most recent event, extensive coral bleaching from hyperthermal stress in the Caribbean occurred in 1997-1998, 2005, and 2010 (Eakin et al., 2010; Heron et al., 2016). If we assume that thermal stress events such as the 2014-2015 event in Florida occur twice per decade up until the estimated timing of annual severe bleaching (2042), we can estimate how many years we have left before local extinction of *D. cylindrus* for each of the percent decline scenarios modeled (80%, 50% and 20%). If the rate of survival throughout the Florida Reef Tract is 80% of colonies, all Florida pillar corals would be lost by 2064. Assuming 50% or 20% survival, we would expect *D. cylindrus* to become extinct in Florida in 2045 and 2031, respectively.

These are simple projections of potential species decline that do not take into account the heterogeneity of abiotic and biotic factors along the Florida Reef Tract. Detailed demographic information about the actual loss of Florida *D. cylindrus* colonies, genotypes, and *S. dendrogyrum* strains are necessary for successful species rehabilitation (Neely et al. in prep). If no active management strategies such as nursery rearing and propagation are enacted, we can expect pillar corals to disappear from Florida within just a few decades. This might be the first reef-building coral species in the Caribbean that will face local if not regional extinction. Global change will continue to alter the composition of reefs if green house gas emissions are not curbed.

### The Consequences of Rare Species Loss

What are the consequences of losing rare species? Experimental removal of rare species was shown to reduce ecosystem resistance to invasion by an exotic grass (Lyons & Schwartz, 2001). It is possible that less common species significantly contribute to the proper maintenance of ecosystem function. Rare plant species were shown to more significantly impact nutrient cycling and retention in an alpine meadow compared to more abundant species (Theodose et al., 1996). In a comprehensive study of species occurrence datasets from coral reefs, alpine meadows, and tropical forests, rare species were repeatedly shown to predominantly support vulnerable functions. These vulnerable functions were defined as ecosystem roles with low redundancy, in that uncommon species with distinct trait combinations bolstered these particular functions (Mouillot et al., 2013). Despite the high diversity in these ecosystems, abundant species did not insure against the services lost by removing rare species. Thus, maintaining these uncommon species is essential to overall community functional diversity.

When two rare species are combined in an obligate symbiosis, then the loss of one species would yield coextinction of the other (Koh et al., 2004). It is possible that *D. cylindrus* sexual recruits can survive by associating with *S. meandrinium* in the absence of *S. dendrogyrum* but the latter has never been found in another coral host species (C. Lewis et al. in prep, A. Lewis et al. in review). It is unknown whether there are free-living strains of *S. dendrogyrum*. Determining the degree to which the association is obligate for the symbiont and the functional roles of the species in the ecosystem is beyond the scope of this study. Without knowing the ecological function of rare marine symbioses such as the one between *D. cylindrus* and *S*. *dendrogyrum*, it is prudent to assume important ecosystem contributions when forming conservation strategies (Lyons et al., 2005).

Continued unprecedented global change will cause further loss of biodiversity. Because species that are both rare and specialized are especially at risk of disappearing (Davies et al., 2004), studies of the ecology and evolution of these species are particularly timely and important. Recent work on *D. cylindrus* (Neely et al. in prep), including this study, has highlighted a lack of recent successful sexual recruitment. The ability of coral species to survive climate change hinges upon ongoing sexual reproduction, enabling selection for more resilient genotypes (Matz et al., 2017). Adaptation is thus extremely unlikely in *D. cylindrus*, necessitating continued *ex situ* efforts to understand spawning, larval development, and larval settlement in this unique Caribbean coral species (Marhaver et al., 2015).

Coral reefs are currently experiencing global decline due to the loss of important herbivores, pollution and nutrient runoff, disease, and climate change (Bruno et al., 2007; Hoegh-Guldberg et al., 2007; Jackson et al., 2014). Since the 1980s, coral cover on Caribbean reefs has already declined by an average of 80% (Gardner et al., 2003). Losing rare species and their corresponding functions may further reduce reef resilience and thus exacerbate coral reef decline beyond what has been predicted (Hoegh-Guldberg et al., 2017; T. P. Hughes et al., 2017).

## Acknowledgements

We thank L Kabay, K Macaulay, L Carne, and the Keys Marine Lab AAUS staff divers for outstanding field support. M Rodriguez-Lanetty (Florida International University) provided some funding and supplies for field sampling. We also thank the staff at the Penn State Genomics Core Facility for running samples as well as M Devlin-Durante and K Lunz for help with obtaining funds. We thank S Kitchen, J Keller, and R Leblois for valuable help with bioinformatics analyses. Computational analyses were conducted on the PSU Institute for CyberScience Advanced CyberInfrastructure high-performance computer cluster. Support for this work was provided by the Florida Fish and Wildlife Conservation Commission’s program, Florida’s Wildlife Legacy Initiative, and the U.S. Fish and Wildlife Service’s State Wildlife Grants program (Marine Projects 2012, T-32). Coral samples were collected under Florida Keys National Marine Sanctuary permits FKNMS-2013-085-A1, FKNMS-2014-004-A1, FKNMS-2016-062, CITES AN001.12US784243/g, and Indigenous Species Research, Retention, and Export permit DFW16005T. Chan was supported by the National Science Foundation (NSF) Graduate Research Fellowship Program under Grant No. DGE1255832. The conclusions are those of the authors and do not necessarily reflect the views of the NSF.

## Data Accessibility

All data affiliated with the manuscript “Fallen Pillars: The Past, Present, and Future Population Dynamics of a Rare, Specialist Coral-Algal Symbiosis” will be made available on Penn State’s publicly accessible repository Scholarsphere (https://scholarsphere.psu.edu/).

*Dendrogyra cylindrus* microsatellite genotypes: available upon article acceptance

*Symbiodinium dendrogyrum* microsatellite genotypes: available upon article acceptance

Raw 454 sequencing data for an adult *Dendrogyra cylindrus* tissue sample from Florida used for microsatellite design: https://scholarsphere.psu.edu/concern/parent/31z40kt37x/file_sets/x6969z327w

3,831 perfect microsatellites with tri- and tetra-nucleotide repeat motif lengths identified using SciRoKo: https://scholarsphere.psu.edu/concern/parent/31z40kt37x/file_sets/tb2773×67v

821 primers designed using Primer3 for the amplification of *Dendrogyra cylindrus* and *Symbiodinium dendrogyrum* microsatellites: https://scholarsphere.psu.edu/concern/parent/31z40kt37x/file_sets/bcc08hg284

## Author Contributions

Andrea N. Chan performed all molecular work and data analysis, and wrote the paper.

Cynthia L. Lewis collected samples and edited the paper.

Karen L. Neely collected samples and edited the paper.

Iliana B. Baums directed the study, obtained funding, provided laboratory space, collected samples, and wrote the paper.

## References

Anderson, D. R. (2007). Model based inference in the life sciences: a primer on evidence: Springer Science & Business Media.

Andras, J. P., Kirk, N. L., Coffroth, M. A., & Harvell, C. (2009). Isolation and characterization of microsatellite loci in *Symbiodinium* B1/B184, the dinoflagellate symbiont of the Caribbean sea fan coral, *Gorgonia ventalina*. Molecular ecology resources, 9(3), 989–993.

Andras, J. P., Kirk, N. L., & Drew Harvell, C. (2011). Range-wide population genetic structure of *Symbiodinium* associated with the Caribbean sea fan coral, *Gorgonia ventalina*. Molecular Ecology, 20(12), 2525–2542.

Andras, J. P., Rypien, K. L., & Harvell, C. D. (2013). Range-wide population genetic structure of the Caribbean sea fan coral, *Gorgonia ventalina*. Molecular Ecology, 22(1), 56–73.

Baums, I. B. (2008). A restoration genetics guide for coral reef conservation. Molecular Ecology, 17(12), 2796–2811.

Baums, I. B., Devlin-Durante, M., Laing, B. A., Feingold, J., Smith, T., Bruckner, A., & Monteiro, J. (2014a). Marginal coral populations: the densest known aggregation of *Pocillopora* in the Galápagos Archipelago is of asexual origin. Frontiers in Marine Science, 1, 59.

Baums, I. B., Devlin-Durante, M. K., & LaJeunesse, T. C. (2014b). New insights into the dynamics between reef corals and their associated dinoflagellate endosymbionts from population genetic studies. Molecular Ecology, 23(17), 4203–4215.

Baums, I. B., Johnson, M., Devlin-Durante, M., & Miller, M. (2010). Host population genetic structure and zooxanthellae diversity of two reef-building coral species along the Florida Reef Tract and wider Caribbean. Coral Reefs, 29(4), 835–842.

Baums, I. B., Miller, M. W., & Hellberg, M. E. (2005). Regionally isolated populations of an imperiled Caribbean coral, *Acropora palmata*. Molecular Ecology, 14(5), 1377–1390.

Baums, I. B., Miller, M. W., & Hellberg, M. E. (2006). Geographic variation in clonal structure in a reef-building Caribbean coral, *Acropora palmata*. Ecological Monographs, 76(4), 503–519.

Besnier, F., & Glover, K. A. (2013). ParallelStructure: A R Package to Distribute Parallel Runs of the Population Genetics Program STRUCTURE on Multi-Core Computers. PLoS ONE, 8(7), e70651. doi:10.1371/journal.pone.0070651

Blanquer, A., & Uriz, M. J. (2010). Population genetics at three spatial scales of a rare sponge living in fragmented habitats. BMC Evolutionary Biology, 10(1), 13.

Boulay, J. N., Hellberg, M. E., Cortés, J., & Baums, I. B. (2014). Unrecognized coral species diversity masks differences in functional ecology. Proceedings of the Royal Society of London B: Biological Sciences, 281(1776), 20131580.

Bruno, J. F., Selig, E. R., Casey, K. S., Page, C. A., Willis, B. L., Harvell, C. D., … Melendy, A. M. (2007). Thermal stress and coral cover as drivers of coral disease outbreaks. PLoS Biology, 5(6), e124.

Budd, A. F. (2000). Diversity and extinction in the Cenozoic history of Caribbean reefs. Coral Reefs, 19(1), 25–35.

Caughley, G. (1994). Directions in conservation biology. Journal of Animal Ecology, 215–244.

Clark-Tapia, R., Mandujano, M. C., Valverde, T., Mendoza, A., & Molina-Freaner, F. (2005). How important is clonal recruitment for population maintenance in rare plant species?: The case of the narrow endemic cactus, *Stenocereus eruca*, in Baja California, México. Biological Conservation, 124(1), 123–132.

Courchamp, F., Clutton-Brock, T., & Grenfell, B. (1999). Inverse density dependence and the Allee effect. Trends in Ecology & Evolution, 14(10), 405–410.

Davies, K. F., Margules, C. R., & Lawrence, J. F. (2004). A synergistic effect puts rare, specialized species at greater risk of extinction. Ecology, 85(1), 265–271.

De Iorio, M., & Griffiths, R. C. (2004a). Importance sampling on coalescent histories. I. Advances in Applied Probability, 36(2), 417–433.

De Iorio, M., & Griffiths, R. C. (2004b). Importance sampling on coalescent histories. II: Subdivided population models. Advances in Applied Probability, 36(2), 434–454.

de Lange, P. J., & Norton, D. A. (2004). The ecology and conservation of *Kunzea sinclairii* (Myrtaceae), a naturally rare plant of rhyolitic rock outcrops. Biological Conservation, 117(1), 49–59.

Devlin-Durante, M. K., & Baums, I. B. (2017). Genome-wide survey of single-nucleotide polymorphisms reveals fine-scale population structure and signs of selection in the threatened Caribbean elkhorn coral, *Acropora palmata*. PeerJ, 5, e4077.

Díaz-Ferguson, E., Haney, R., Wares, J., & Silliman, B. (2010). Population genetics of a trochid gastropod broadens picture of Caribbean Sea connectivity. PloS one, 5(9), e12675.

Dorken, M. E., & Eckert, C. G. (2001). Severely reduced sexual reproduction in northern populations of a clonal plant, *Decodon verticillatus* (Lythraceae). Journal of Ecology, 89(3), 339–350.

Eakin, C. M., Morgan, J. A., Heron, S. F., Smith, T. B., Liu, G., Alvarez-Filip, L., … Bouchon, C. (2010). Caribbean corals in crisis: record thermal stress, bleaching, and mortality in 2005. PloS one, 5(11), e13969.

Earl, D. A. (2012). STRUCTURE HARVESTER: a website and program for visualizing STRUCTURE output and implementing the Evanno method. Conservation Genetics Resources, 4(2), 359–361.

Edmunds, P., Roberts, D., & Singer, R. (1990). Reefs of the northeastern Caribbean I. Scleractinian populations. Bulletin of Marine Science, 46(3), 780–789.

Ehrenburg, C. (1834). Beitrage zur physiologischen Kenntniss der Corallenthiere im allgemeined, und besonders des rothen Meers, nebst einem Versucch zu physiologishen systematic derselben. Phys Abh Konigl Akad Wissech Berlin aus der Jahar, 1832, 225–380.

Ellstrand, N. C., & Elam, D. R. (1993). Population genetic consequences of small population size: implications for plant conservation. Annual Review of Ecology and Systematics, 24(1), 217–242.

Estoup, A., & Angers, B. (1998). Microsatellites and minisatellites for molecular ecology: theoretical and empirical considerations. Advances in Molecular Ecology. Edited by: Carvalho GR.: Amsterdam: IOS Press.

Evanno, G., Regnaut, S., & Goudet, J. (2005). Detecting the number of clusters of individuals using the software STRUCTURE: a simulation study. Molecular Ecology, 14(8), 2611–2620.

Excoffier, L., Smouse, P. E., & Quattro, J. M. (1992). Analysis of molecular variance inferred from metric distances among DNA haplotypes: application to human mitochondrial DNA restriction data. Genetics, 131(2), 479–491.

Finney, J. C., Pettay, D. T., Sampayo, E. M., Warner, M. E., Oxenford, H. A., & LaJeunesse, T. C. (2010). The relative significance of host–habitat, depth, and geography on the ecology, endemism, and speciation of coral endosymbionts in the genus *Symbiodinium*. Microbial Ecology, 60(1), 250–263.

Flather, C. H., & Sieg, C. H. (2007). Species rarity: definition, causes and classification. Conservation of rare or little-known species: Biological, social, and economic considerations, 40–66.

Foster, N. L., Paris, C. B., Kool, J. T., Baums, I. B., Stevens, J. R., Sanchez, J. A., … Day, O. (2012). Connectivity of Caribbean coral populations: complementary insights from empirical and modelled gene flow. Molecular Ecology, 21(5), 1143–1157.

Gardner, T. A., Côté, I. M., Gill, J. A., Grant, A., & Watkinson, A. R. (2003). Long-term region-wide declines in Caribbean corals. Science, 301(5635), 958–960.

Goreau, T. F., & Wells, J. (1967). The shallow-water Scleractinia of Jamaica: revised list of species and their vertical distribution range. Bulletin of Marine Science, 17(2), 442–453.

Hedrick, P. W. (2005). A standardized genetic differentiation measure. Evolution, 59(8), 1633–1638.

Heron, S. F., Maynard, J. A., Van Hooidonk, R., & Eakin, C. M. (2016). Warming trends and bleaching stress of the world’s coral reefs 1985–2012. Scientific reports, 6, 38402.

Hoegh-Guldberg, O., Mumby, P. J., Hooten, A. J., Steneck, R. S., Greenfield, P., Gomez, E., … Caldeira, K. (2007). Coral reefs under rapid climate change and ocean acidification. Science, 318(5857), 1737–1742.

Hoegh-Guldberg, O., Poloczanska, E. S., Skirving, W., & Dove, S. (2017). Coral reef ecosystems under climate change and ocean acidification. Frontiers in Marine Science, 4, 158.

Hughes, A. R., & Stachowicz, J. J. (2004). Genetic diversity enhances the resistance of a seagrass ecosystem to disturbance. Proceedings of the National Academy of Sciences of the United States of America, 101(24), 8998–9002.

Hughes, T. P., Barnes, M. L., Bellwood, D. R., Cinner, J. E., Cumming, G. S., Jackson, J. B., … Morrison, T. H. (2017). Coral reefs in the Anthropocene. Nature, 546(7656), 82.

Hunter, I., & Jones, B. (1996). Coral associations of the Pleistocene Ironshore Formation, Grand Cayman. Coral Reefs, 15(4), 249–267.

Jackson, J., Donovan, M., Cramer, K., & Lam, V. (2014). Status and trends of Caribbean coral reefs: 1970-2012. Retrieved from

Jossart, Q., De Ridder, C., Lessios, H. A., Bauwens, M., Motreuil, S., Rigaud, T., … David, B. (2017). Highly contrasted population genetic structures in a host–parasite pair in the Caribbean Sea. Ecology and evolution, 7(22), 9267–9280.

Jost, L. (2008). GST and its relatives do not measure differentiation. Molecular Ecology, 17(18), 4015–4026.

Kabay, L. (2016). Population Demographics and Sexual Reproduction Potential of the Pillar Coral, Dendrogyra cylindrus, on the Florida Reef Tract.

Kemp, D., Fitt, W., & Schmidt, G. (2008). A microsampling method for genotyping coral symbionts. Coral Reefs, 27(2), 289–293.

Kofler, R., Schlötterer, C., & Lelley, T. (2007). SciRoKo: a new tool for whole genome microsatellite search and investigation. Bioinformatics, 23(13), 1683–1685.

Koh, L. P., Dunn, R. R., Sodhi, N. S., Colwell, R. K., Proctor, H. C., & Smith, V. S. (2004). Species coextinctions and the biodiversity crisis. Science, 305(5690), 1632–1634.

Kopelman, N. M., Mayzel, J., Jakobsson, M., Rosenberg, N. A., & Mayrose, I. (2015). Clumpak: a program for identifying clustering modes and packaging population structure inferences across K. Molecular ecology resources, 15(5), 1179–1191.

Leblois, R., Pudlo, P., Néron, J., Bertaux, F., Reddy Beeravolu, C., Vitalis, R., & Rousset, F. (2014). Maximum-likelihood inference of population size contractions from microsatellite data. Molecular Biology and Evolution, 31(10), 2805–2823.

Lewis, A. M., Chan, A. N., & LaJeunesse, T. C. (2018). New species of endosymbiotic dinoflagellates specific to common reef-building corals across the Greater Caribbean. Journal of Eukaryotic Microbiology, In Review.

Lewis, C. L., Neely, K. L., Richardson, L. L., & Rodriguez-Lanetty, M. (2017). Temporal dynamics of black band disease affecting pillar coral (*Dendrogyra cylindrus*) following two consecutive hyperthermal events on the Florida Reef Tract. Coral Reefs, 36(2), 427–431.

Lunz, K., Neely, K. L., Kabay, L., & Gilliam, D. (2016). Monitoring and Mapping of Dendrogyra cylindrus of the Florida Reef Tract: Final Report (9751-281-1168). Retrieved from

Lyons, K. G., Brigham, C., Traut, B., & Schwartz, M. W. (2005). Rare species and ecosystem functioning. Conservation Biology, 19(4), 1019–1024.

Lyons, K. G., & Schwartz, M. W. (2001). Rare species loss alters ecosystem function–invasion resistance. Ecology Letters, 4(4), 358–365.

Manzello, D. P. (2015). Rapid recent warming of coral reefs in the Florida Keys. Scientific reports, 5, 16762.

Marhaver, K. L., Vermeij, M. J., & Medina, M. M. (2015). Reproductive natural history and successful juvenile propagation of the threatened Caribbean Pillar Coral *Dendrogyra cylindrus*. BMC Ecology, 15(1), 1. doi:http://dx.doi.org/10.1186/s12898-015-0039-7

Matz, M. V., Treml, E. A., Aglyamova, G. V., van Oppen, M. J., & Bay, L. K. (2017). Potential for rapid genetic adaptation to warming in a Great Barrier Reef coral. bioRxiv.

McKinney, M. L. (1997). Extinction vulnerability and selectivity: combining ecological and paleontological views. Annual Review of Ecology and Systematics, 28(1), 495–516.

Meirmans, P. G., & Van Tienderen, P. H. (2004). GENOTYPE and GENODIVE: two programs for the analysis of genetic diversity of asexual organisms. Molecular Ecology Notes, 4(4), 792–794.

Miller, M., Baums, I., Pausch, R., Bright, A., Cameron, C., Williams, D., … Woodley, C. (2018). Clonal structure and variable fertilization success in Florida Keys broadcast-spawning corals. Coral Reefs, 37(1), 239–249.

Miller, S., Precht, W. F., Rutten, L. M., & Chiappone, M. (2013). Florida Keys population abundance estimates for nine coral species proposed for listing under the US Endangered Species Act.

Miller, S. L., Chiappone, M., & Rutten, L. M. (2010). Abundance, Distribution and Condition of Benthic Coral Reef Organisms in the Upper Florida Keys National Marine Sanctuary: 2010 Quick Look Report and Data Summary: University of North Carolina at Wilmington.

Mouillot, D., Bellwood, D. R., Baraloto, C., Chave, J., Galzin, R., Harmelin-Vivien, M., … Mouquet, N. (2013). Rare species support vulnerable functions in high-diversity ecosystems. PLoS Biology, 11(5), e1001569.

Nei, M. (1987). Molecular evolutionary genetics: Columbia university press.

NOAA. (2014). Endangered and threatened wildlife and plants: final listing determinations on proposal to list 66 reef-building coral species and to reclassify elkhorn and staghorn corals. Fed Reg(75), 271.

Parkinson, J. E., Banaszak, A. T., Altman, N. S., LaJeunesse, T. C., & Baums, I. B. (2015). Intraspecific diversity among partners drives functional variation in coral symbioses. Scientific reports, 5, 15667.

Parkinson, J. E., & Baums, I. B. (2014). The extended phenotypes of marine symbioses: Ecological and evolutionary consequences of intraspecific genetic diversity in coral–algal associations. Frontiers in microbiology, 5, 445.

Peakall, R., & Smouse, P. E. (2006). GENALEX 6: genetic analysis in Excel. Population genetic software for teaching and research. Molecular Ecology Notes, 6(1), 288–295.

Pettay, D. T., & LaJeunesse, T. C. (2007). Microsatellites from clade B *Symbiodinium* spp. specialized for Caribbean corals in the genus *Madracis*. Molecular Ecology Notes, 7(6), 1271–1274.

Precht, W. F., Gintert, B. E., Robbart, M. L., Fura, R., & Van Woesik, R. (2016). Unprecedented disease-related coral mortality in Southeastern Florida. Scientific reports, 6, 31374.

Pritchard, J. K., Stephens, M., & Donnelly, P. (2000). Inference of population structure using multilocus genotype data. Genetics, 155(2), 945–959.

Pritchard, J. K., Wen, X., & Falush, D. (2009). Documentation for structure software: Version 2.3.

Puechmaille, S. J. (2016). The program structure does not reliably recover the correct population structure when sampling is uneven: subsampling and new estimators alleviate the problem. Molecular ecology resources, 16(3), 608–627.

Rabinowitz, D. (1981). Seven forms of rarity. The biological aspects of rare plants conservation, 205–217.

Rabinowitz, D., Rapp, J. K., & Dixon, P. M. (1984). Competitive abilities of sparse grass species: means of persistence or cause of abundance. Ecology, 65(4), 1144–1154.

Reusch, T. B., Ehlers, A., Hämmerli, A., & Worm, B. (2005). Ecosystem recovery after climatic extremes enhanced by genotypic diversity. Proceedings of the National Academy of Sciences of the United States of America, 102(8), 2826–2831.

Rippe, J. P., Matz, M. V., Green, E. A., Medina, M., Khawaja, N. Z., Pongwarin, T., … Davies, S. W. (2017). Population structure and connectivity of the mountainous star coral, *Orbicella faveolata*, throughout the wider Caribbean region. Ecology and evolution, 7(22), 9234–9246.

Rousset, F. (2008). genepop’007: a complete re-implementation of the genepop software for Windows and Linux. Molecular ecology resources, 8(1), 103–106.

Rozen, S., & Skaletsky, H. (1999). Primer3 on the WWW for general users and for biologist programmers. Bioinformatics methods and protocols, 365–386.

Ryman, N., & Palm, S. (2006). POWSIM: a computer program for assessing statistical power when testing for genetic differentiation. Molecular Ecology Notes, 6(3), 600–602.

Santos, S. R., & Coffroth, M. A. (2003). Molecular genetic evidence that dinoflagellates belonging to the genus *Symbiodinium* Freudenthal are haploid. The Biological Bulletin, 204(1), 10–20.

Scobie, A., & Wilcock, C. (2009). Limited mate availability decreases reproductive success of fragmented populations of *Linnaea borealis*, a rare, clonal self-incompatible plant. Annals of Botany, 103(6), 835–846.

Shoguchi, E., Shinzato, C., Kawashima, T., Gyoja, F., Mungpakdee, S., Koyanagi, R., … Fujiwara, M. (2013). Draft assembly of the *Symbiodinium minutum* nuclear genome reveals dinoflagellate gene structure. Current Biology, 23(15), 1399–1408.

Silvertown, J. (2008). The evolutionary maintenance of sexual reproduction: evidence from the ecological distribution of asexual reproduction in clonal plants. International Journal of Plant Sciences, 169(1), 157–168.

Steiner, S. C., & Kerr, J. (2008). Stony corals in Dominica during the 2005 bleaching episode and one year later. Revista de Biologia Tropical, 56(1).

Stephens, P. A., & Sutherland, W. J. (1999). Consequences of the Allee effect for behaviour, ecology and conservation. Trends in Ecology & Evolution, 14(10), 401–405.

Stocker, T., Qin, D., Plattner, G., Tignor, M., Allen, S., Boschung, J., … Midgley, B. (2013). IPCC, 2013: climate change 2013: the physical science basis. Contribution of working group I to the fifth assessment report of the intergovernmental panel on climate change.

Swarts, N. D., Sinclair, E. A., Francis, A., & Dixon, K. W. (2010). Ecological specialization in mycorrhizal symbiosis leads to rarity in an endangered orchid. Molecular Ecology, 19(15), 3226–3242.

Szmant, A. M. (1986). Reproductive ecology of Caribbean reef corals. Coral Reefs, 5(1), 43–53. doi:http://dx.doi.org/10.1007/bf00302170

Theodose, T. A., Jaeger III, C. H., Bowman, W. D., & Schardt, J. C. (1996). Uptake and allocation of 15 N in alpine plants: implications for the importance of competitive ability in predicting community structure in a stressful environment. Oikos, 59–66.

Thornhill, D., Howells, E., Wham, D., Steury, T., & Santos, S. (2017). Population genetics of reef coral endosymbionts (*Symbiodinium*, Dinophyceae). Molecular Ecology, 26(10), 2640–2659.

Tooker, J. F., & Frank, S. D. (2012). Genotypically diverse cultivar mixtures for insect pest management and increased crop yields. Journal of Applied Ecology, 49(5), 974–985.

van Hooidonk, R., Maynard, J., Tamelander, J., Gove, J., Ahmadia, G., Raymundo, L., … Planes, S. (2017). Coral Bleaching Futures: Downscaled Projections of Bleaching Conditions for the World’s Coral Reefs, Implications of Climate Policy and Management Responses.

van Hooidonk, R., Maynard, J. A., Liu, Y., & Lee, S. K. (2015). Downscaled projections of Caribbean coral bleaching that can inform conservation planning. Global Change Biology, 21(9), 3389–3401.

Vera, J. C., Wheat, C. W., Fescemyer, H. W., Frilander, M. J., Crawford, D. L., Hanski, I., & Marden, J. H. (2008). Rapid transcriptome characterization for a nonmodel organism using 454 pyrosequencing. Molecular Ecology, 17(7), 1636–1647.

Vollmer, S. V., & Palumbi, S. R. (2006). Restricted gene flow in the Caribbean staghorn coral *Acropora cervicornis*: implications for the recovery of endangered reefs. Journal of Heredity, 98(1), 40–50.

Waits, L. P., Luikart, G., & Taberlet, P. (2001). Estimating the probability of identity among genotypes in natural populations: cautions and guidelines. Molecular Ecology, 10(1), 249–256.

Ward, J., Rypien, K., Bruno, J., Harvell, C., Jordan-Dahlgren, E., Mullen, K., … Smith, G. (2006). Coral diversity and disease in Mexico. Diseases of Aquatic Organisms, 69(1), 23–31.

Warner, R. R., & Chesson, P. L. (1985). Coexistence mediated by recruitment fluctuations: a field guide to the storage effect. The American Naturalist, 125(6), 769–787.

Werth, S., & Scheidegger, C. (2012). Congruent genetic structure in the lichen-forming fungus *Lobaria pulmonaria* and its green-algal photobiont. Molecular Plant-Microbe Interactions, 25(2), 220–230.

Willing, E.-M., Dreyer, C., & Van Oosterhout, C. (2012). Estimates of genetic differentiation measured by FST do not necessarily require large sample sizes when using many SNP markers. PloS one, 7(8), e42649.

Wright, S. (1951). The Genetical Structure of Populations. Annals of Eugenics, 15, 323–354.

